# Retained introns in long RNA-seq reads are not reliably detected in sample-matched short reads

**DOI:** 10.1101/2022.03.11.484016

**Authors:** Julianne K. David, Sean K. Maden, Mary A. Wood, Reid F. Thompson, Abhinav Nellore

## Abstract

There is growing interest in retained introns in a variety of disease contexts including cancer and aging. Many software tools have been developed to detect retained introns from short RNA-seq reads, but reliable detection is complicated by overlapping genes and transcripts as well as the presence of unprocessed or partially processed RNAs. We compared introns detected by 5 tools using short RNA-seq reads with introns observed in long RNA-seq reads from the same biological specimens and found: (1) significant disagreement among tools (Fleiss’ ***κ* = 0.231**) such that **52.4%** of all detected intron retentions were not called by more than one tool; (2) that no tool achieved greater than **20%** precision or **35%** recall under generous conditions; and (3) that retained intron detectability was adversely affected by greater intron length and overlap with annotated exons.

## 1 Introduction

During RNA transcription, multiple spliceosomes may act on the same transcript in parallel to remove segments of sequence called introns and splice together flanking exons [1]. Most splicing occurs stochastically [2] during transcription [3–5], although up to 20% of splicing may occur after transcription and polyadenylation [5, 6] (Fig. S1). Introns are spliced by several known spliceosome types, of which the most-studied are called U2 and U12 [7]. Splicing is known to occur primarily in the nucleus [8], though there is evidence of cytoplasmic splicing [9–12].

Intron retention (IR) is a form of alternative splicing where an intron normally spliced out during transcript processing remains after processing is complete. IR occurs in up to 80% of protein-coding genes in humans [13] and may affect gene expression regulation [14– 20] as well as response to stress [21–23]. Transcripts containing introns may also be stably detained in the nucleus before undergoing delayed splicing (“intron detention,” or ID), with implications for temporal gene expression [24]. In cancers, high levels of IR [25–27] can generate aberrant splicing products with known and potential biological consequences for gene expression and cell survival [28]. IR rarely gives rise to a protein product [29, 30], but novel peptides derived from transcripts with retained introns (RIs) are increasingly being studied in disease contexts such as cancer [31–35].

Despite its biological relevance, detection of IR from bulk RNA sequencing (RNA-seq) data remains challenging for two principal reasons: (1) A short RNA-seq read (e.g., from Illumina’s HiSeq, NovaSeq, or MiSeq platforms) is almost never long enough to resolve a full intron or its context in a transcript, particularly in genome regions with multiple overlapping transcripts; (2) RNA-seq data may contain intronic sequence from unprocessed or partially processed transcripts, DNA contamination, and non-messenger RNA such as circular RNAs (cRNAs) [4, 36], potentially yielding spurious IR calls, independent of read length.

Existing tools designed specifically for RI detection make simplifying assumptions to address the above issues. These tools include keep me around (KMA) [37], IntEREst [38], iREAD [39], superintronic [40], and IRFinder [13] and its most recent implementation as IRFinder-S [41]. Some mitigate challenge (1) by ignoring from consideration any intronic regions that overlap other features (KMA, IntEREst, iREAD), leaving biological blindspots in RI detection [37–39]. Some attempt to mitigate challenge (2) by recommending that a user provides poly(A)-selected data as their input [13, 37, 39, 40], assuming that poly(A) selected data represents fully processed, mature RNA. However, poly(A) selection during library preparation has been shown not to remove all immature post-transcriptionally spliced RNA molecules, and intronic sequences are commonly found in poly(A)-selected RNA-sequencing data [42, 43]. To clarify the quality of and best practices for RI detection, we performed tests on poly(A)-selected, sample-matched long- and short-read sequencing runs for two biological specimens, with processed long-read data providing a standard against which we evaluated short read-based RI detection.

## 2 Results

### 2.1 Testing RI detection using sample-paired short- and deep long-read RNA-seq data

To generate a dataset to test RI detection, we identified two human biological specimens on the Sequence Read Archive (SRA) with RNA-seq data from both Illumina short-read and PacBio Iso-Seq RS II long-read platforms (Fig. 1). These were a human whole blood sample (HX1) [44] and a human induced pluripotent stem cell line sample (iPSC) [45], with, respectively, 46 and 27 Iso-Seq runs, 24.4 and 91.3 million aligned short reads, and 945 and 840 thousand aligned long reads (Table S1). To confine attention to robustly represented loci, we identified a set of 4,369 and 4,639 target genes in HX1 and iPSC samples, respectively, each with ≥ 2 short reads per base median coverage across the full gene length and ≥ 5 long reads assigned to at least one isoform of the gene (Fig. S2).

**Fig. 1:**
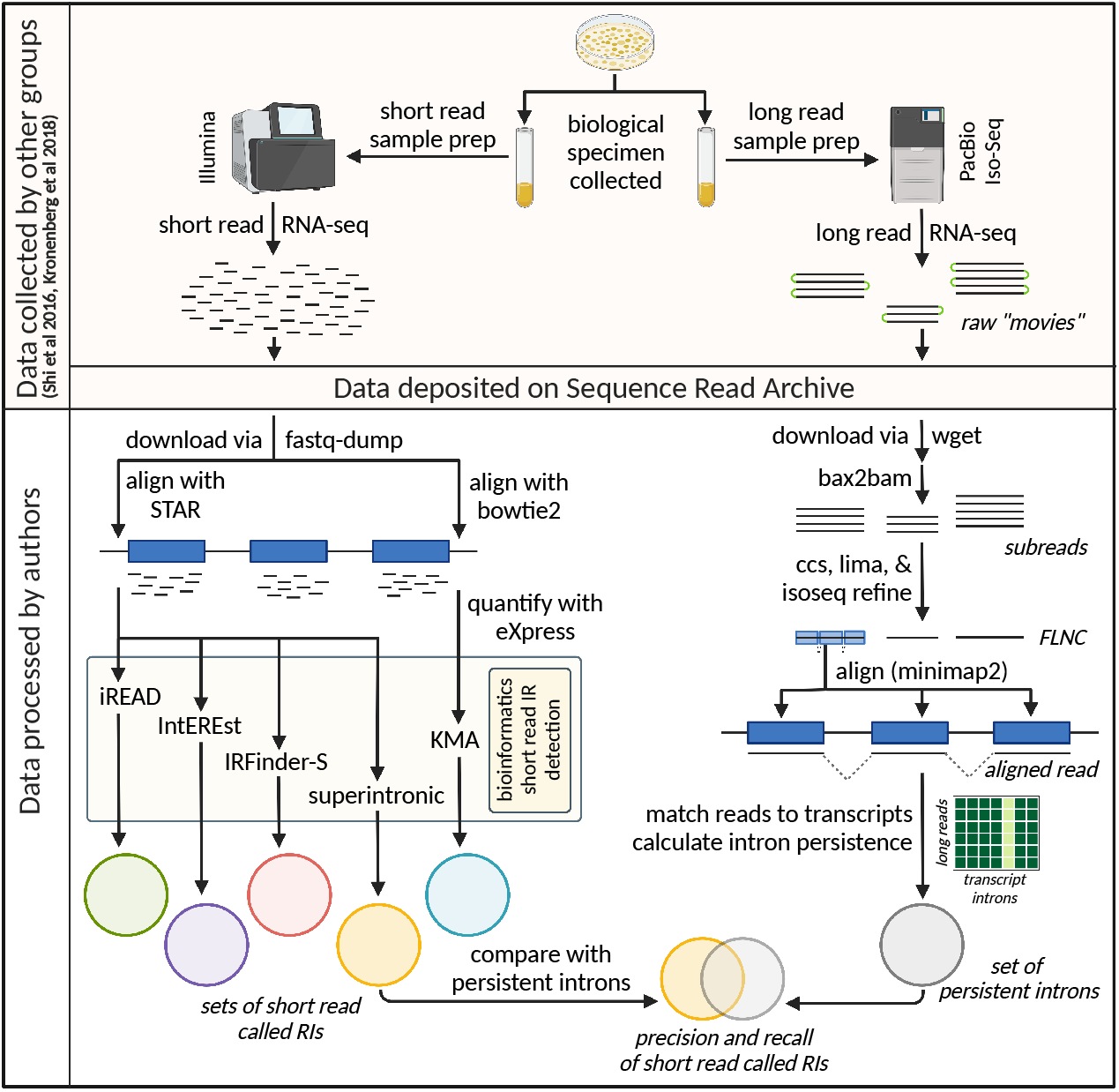
Overview of experimental plan. Long and short read RNA-seq data from the same biological specimen [44, 45] were downloaded from the SRA and subject to processing and analysis. Short reads (left path) were aligned and quantified according to the requirements of five short read RI detection tools [37–41], and retained introns were called by each of these. The raw long Iso-Seq reads (right path) were processed to the stage of full-length non-concatemer (FLNC) reads, but left unclustered. After long reads were aligned to the reference genome, each aligned read was assigned to a best match transcript or discarded, and intron persistence was calculated. The called RI output of each short read detection tool was compared against the set of persistent introns identified in the long read data (where *P_i_* ≥ 0.1). Created with BioRender.com.

We sought to quantify IR in each biological specimen using long-read data, accounting for random splicing and sample contamination that may lead to noisy splicing patterns. For a given intron *i* and transcript *t*, we defined persistence *P_i,t_* as

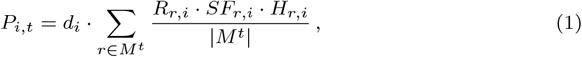

where *r* is a read among the set of all reads *M ^t^* assigned as best matches to transcript *t*, information density *d_i_* is the proportion of *M ^t^* covering intron *i*, the binary variable *R_r,i_* is 1 if and only if *r* provides evidence for the retention of *i*, and the spliced fraction *SF_r,i_* and scaled Hamming similarity *H_r,i_* are defined in Methods (see Equations 3 and 4). In brief, the intron persistence *P_i,t_* incorporates the extent and similarity of splicing across transcript reads, accounting for stochastic splicing initiation and progression (Fig. S1). Finally, to address ambiguity in transcripts of origin in short-read data, we defined intron *i*’s persistence *P_i_* as the maximum persistence across all isoforms *T_i_* that contain *i*:

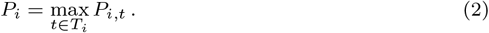

Going forward, we define a “persistent intron” as an intron for which *P_i_* ≥ 0.1.

Across all transcripts studied in both samples, a substantial majority (83.7%) of introns were fully spliced out (*P_i,t_* = 0), and a small minority (0.15%) of introns were always unspliced within a transcript (*P_i,t_* = 1) (Figs. 2a and S3). These extreme values are in keeping with our qualitative understanding of splicing patterns; however, the range of intermediate persistence values appears to represent a spectrum with varying extents of inconsistent splicing across and between reads. While we tested short-read RI detection on a per-sample basis, we also compared intron persistence patterns between HX1 and iPSC samples and found significant similarity in splicing patterns across matched transcripts (Figs. S3 and S4).

**Fig. 2:**
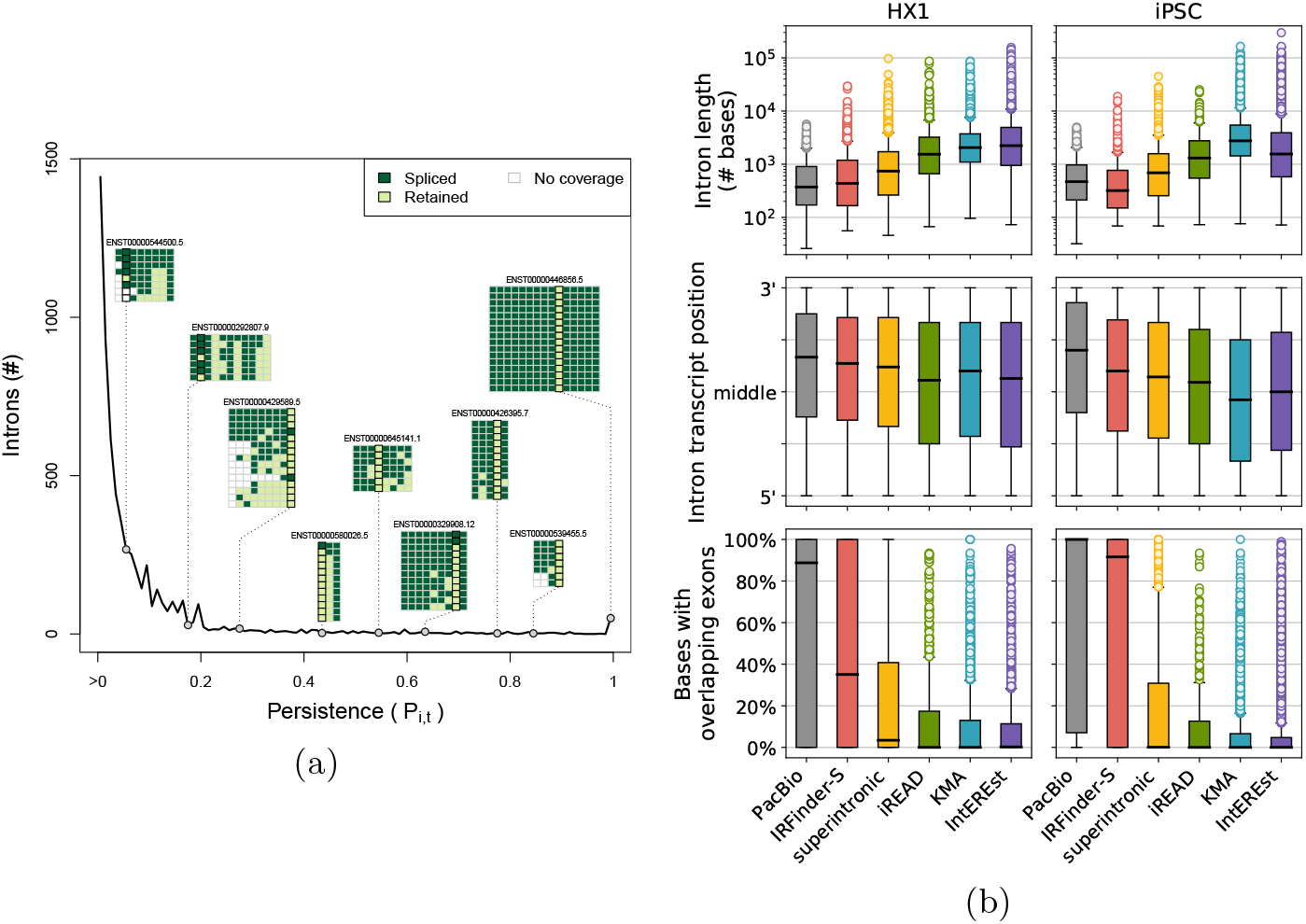
(a) Distribution of persistence *P_i,t_* and representative transcript examples for iPSC. The number of introns (y-axis) having a given persistence value (x-axis) is shown as a dark black line; note that a large number of introns with *P_i_* = 0 are omitted from this analysis. Along the line, gray circles indicate the *P_i_* value corresponding to each of nine introns from representative transcript examples (each transcript is labeled by Ensembl ID, e.g., ENST00000446856.5). Read-level data is shown for each transcript as a colored matrix, where each row is a single long read assigned to the transcript and each column represents a given intron, and color indicates whether an intron is retained (light green), spliced out (dark green), or lacking sequence coverage (white) in a given read. (b) Distributions of properties of persistent and called RIs. Each panel contains a series of boxplots depicting the distribution of intron length (top, log-scale), relative position in transcript (middle), and % of intron bases with overlapping annotated exons (bottom) for HX1 (left) and iPSC (right). The distribution of each of these features is shown for long-read persistent introns (“PacBio”, gray) and RIs called by each of the five short read tools: IRFinder-S (red), superintronic (yellow), iREAD (green), KMA (blue), IntEREst (purple).

### 2.2 Similarities of intron properties across short-read RI detection tool outputs

We compared RIs called by five tools for short-read data (Table 1). While most introns were consistently spliced out, 39.9% (1,743/4,369) and 31.4% (1,457/4,639) of target genes in HX1 and iPSC, respectively, had at least one RI identified in either short- or long-read data. Expression of called RIs varied substantially between tools in both HX1 (Fleiss’ *κ* = 0.282) and iPSC (Fleiss’ *κ* = 0.162), though we did observe moderate overall correlation between the output of IntEREst, superintronic, and KMA (Fig. S5). Further, using circBase [46] to probe whether cRNA contamination may have affected RI detection, we identified only a small percent (< 5%) of called RIs that appeared to overlap intronic cRNAs (Fig. S6).

**Table 1:**
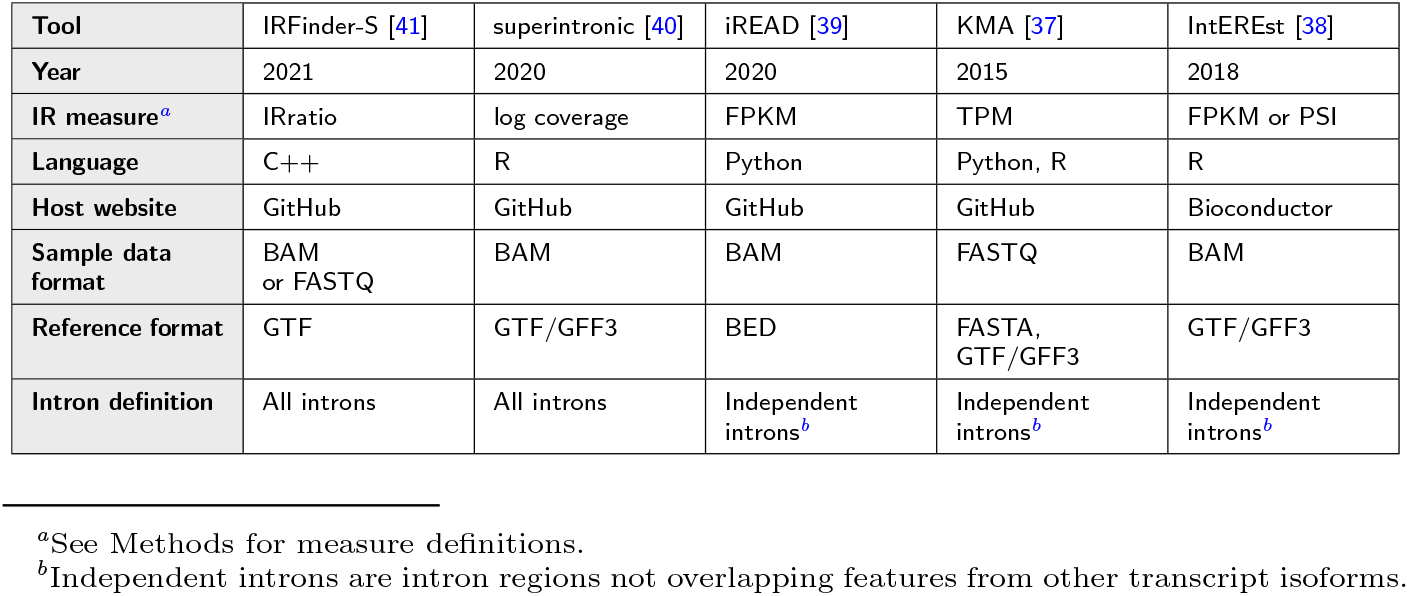
Short-read tools studied.

We next examined the distributions of several intron properties (length, relative position in transcript, and annotated exon overlap) and their relationships with the set of RIs called by each short-read tool and their relative expression levels (Figs. 2b and S7). Unsurprisingly, tools that exclude introns with overlapping genomic features (i.e. KMA, IntEREst, iREAD; Table 1) had exceedingly low overlap between exons and the IRs they reported. We also note that KMA and IntEREst called extremely long RIs (up to > 297 kilobases), compared to those called by other short-read tools or the persistent introns identified from long read data (maximum 6,275 and 5,926 bases in HX1 and iPSC). We observed a slight overall 3’ bias among persistent introns from long-read data, as well as the set of RIs from several short-read tools (Fig. 2b), potentially reflecting the relatively shorter duration of exposure of 3’ introns to the cotranscriptional splicing machinery and/or implicit 3’ bias of the Clontech sample prep [47] used in both samples [44, 45]. Despite this slight 3’ tendency, there was no appreciable association between intron persistence and intron position in transcript (Fig. S8). Among all tools, IRFinder-S called a set of RIs with characteristics most similar to persistent introns from long-read data (Fig. 2b).

### 2.3 Precision and recall are poor across short-read RI detection tools

We tested performance (precision, recall, and F1-score) of RI detection by five short-read tools, comparing sets of called RIs against persistent introns identified from long read data (defined as *P_i_* ≥ 0.1). Overall tool performance was poor in all cases (Fig. 3a, Table S2). Many persistent introns (55% and 48% in iPSC and HX1, respectively, Fig. S9) were not called by any short-read tool, and the majority of called RIs were neither identified among persistent introns in long-read data nor consistently called between short-read tools (Figs. 3c and S9). In HX1 and iPSC, respectively, 54% and 49% of called RIs were not called by more than one tool (52.4% overall). IRFinder-S had the best performance across most metrics, possibly due to the similarity between the properties of its called RIs and properties of persistent introns. By contrast, iREAD demonstrated the lowest recall across all tools, likely due to its sparse calling of RIs (Fig. S10). Performance metrics for IntEREst and KMA were very similar across both samples (Fig. 3b).

**Fig. 3:**
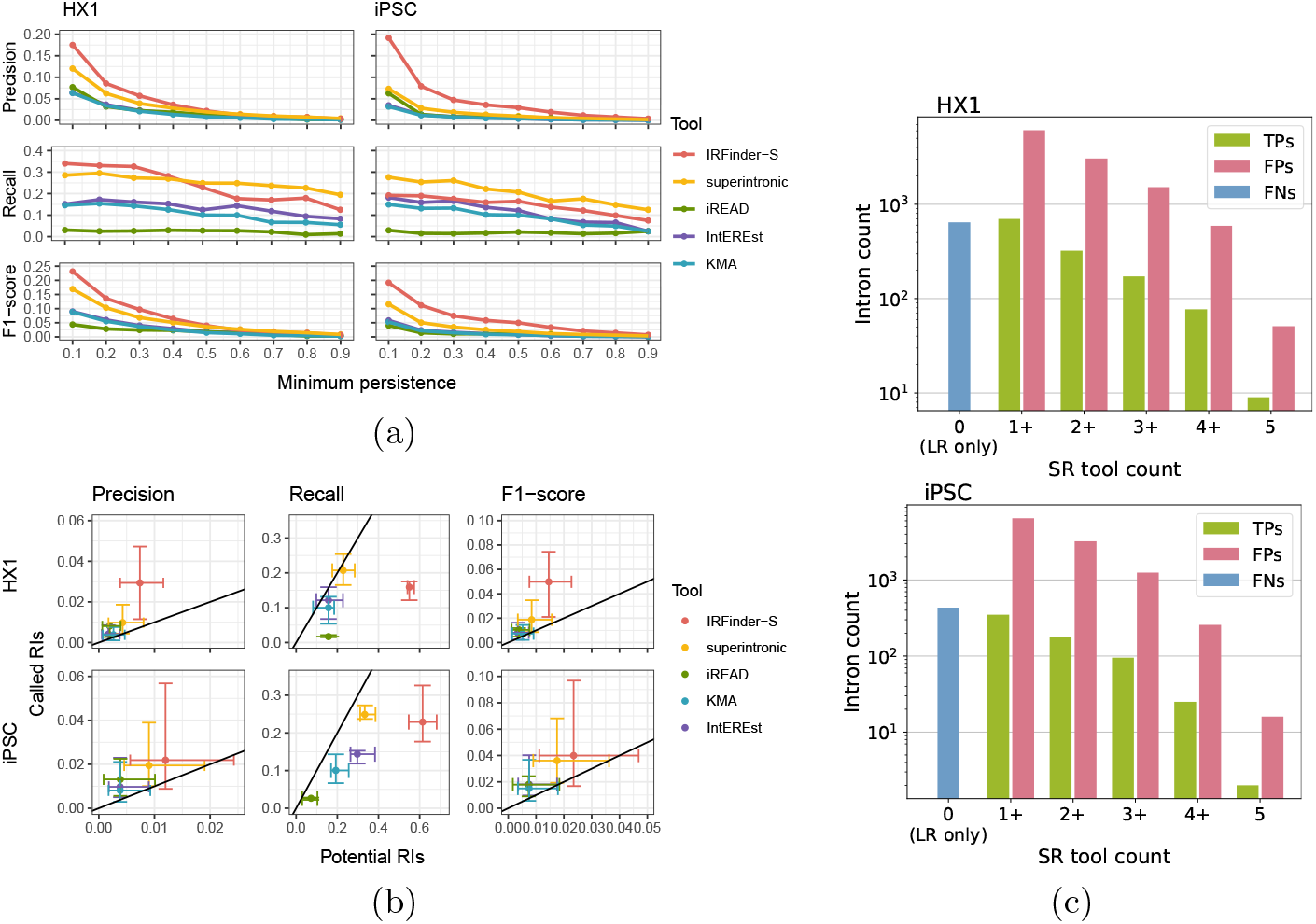
(a) Short-read tool performance across different thresholds of intron persistence. Each panel displays tool performance along the y-axis (measured by one of precision, recall, or F1-score as labeled) for a set of introns defined by the indicated threshold for intron persistence along the x-axis. Data for HX1 and iPSC are shown at left and right, respectively, with each tool’s per-sample performance depicted in a different color (IRFinder-S [red], superintronic [yellow], iREAD [green], IntEREst [purple], and KMA [blue]). (b) Variation in short-read tool performance across intron persistence thresholds for potential vs. called RIs. Each panel displays tool performance as measured by precision (left), recall (middle), and F1-score (right) for HX1 (top) and iPSC (bottom) samples. The performances for each tool’s potential RIs and called RIs are shown along the x- and y-axes, respectively, with centroid and whiskers denoting, respectively, the median and interquartile range of tool performance across intron persistence thresholds. Each tool’s performance is depicted in a different color (IRFinder-S [red], superintronic [yellow], iREAD [green], IntEREst [purple], and KMA [blue]). Reference lines are shown with slope of 1. (c) Varying degrees of consensus of retained intron calls among short-read tools. Bar plots depict the number of true positive (green), false positive (pink), and false negative (blue) intron calls (y-axis) consistent across a specified number of short-read (SR) tools (x-axis). Upper and lower panels depict HX1 and iPSC data, respectively. LR denotes long-read data.

To address sensitivity in persistent intron identification, we also considered short-read tool performance on subsets of long-read introns with increasing minimum thresholds of intron persistence (*P_i_* ≥ 0.1 to 0.9 in increments of 0.1). We found that overall performance remained poor across all levels of intron persistence, with uniformly worse precision, recall and F1-score as intron persistence increased (Figs. 3a and S11). While individual tool performance varied significantly, IRFinder-S and superintronic were consistently best performers, albeit interchangeably depending on the sample, metric assessed, and intron persistence threshold. For instance, IRFinder-S demonstrated highest recall in HX1 at the lowest cut-off values (*P_i_* ≥ 0.1 to 0.4), while superintronic demonstrated higher recall across higher thresholds in HX1 and for all cutoffs in iPSC (Table S2).

Finally, since each tool is capable of calling RIs with different levels of stringency, we evaluated tool performance on a raw set of all potential RIs (all expressed introns detected by that tool) vs. the corresponding subset of introns called as RIs by that tool. Rather than improving overall performance by retaining persistent RIs and removing false positive ones, stringency filters improved precision at the expense of recall, with a slight corresponding improvement in F1-score across tools (Fig. 3b, Supplementary File [`supplementary_performance_data.xlsx`]).

### 2.4 Short introns and introns that do not overlap exons are more reliably called

We next compared the distributions of six intronic properties (length, position, exonic overlap, splice site motifs, U2- vs. U12-type spliceosomes, and uniformity of coverage by mapped reads) between the sets of true positive (TP), false positive (FP) and false negative (FN) RIs for each tool. Every tool except IRFinder-S had difficulty identifying shorter RIs (< 600 bases) (Figs. 4a and 4b). FPs tended to be longer than either TPs or FNs, and were distributed more centrally within a transcript compared to persistent introns (both TPs and FNs) across all tools (Figs. 4a and S12). Further, there was a relative 3’ bias for the small subset of FPs that were shared across all short-read tools, potentially reflective of the minimum coverage filters for most tools combined with sequencing coverage bias [48] (Fig. S13). As expected, the overwhelming majority of introns across all tools had canonical GT-AG splice motifs and splicing by the U2 spliceosome, while FNs showed increased frequencies of other motifs and spliceosome types relative to FPs and TPs (Fig. S14).

**Fig. 4:**
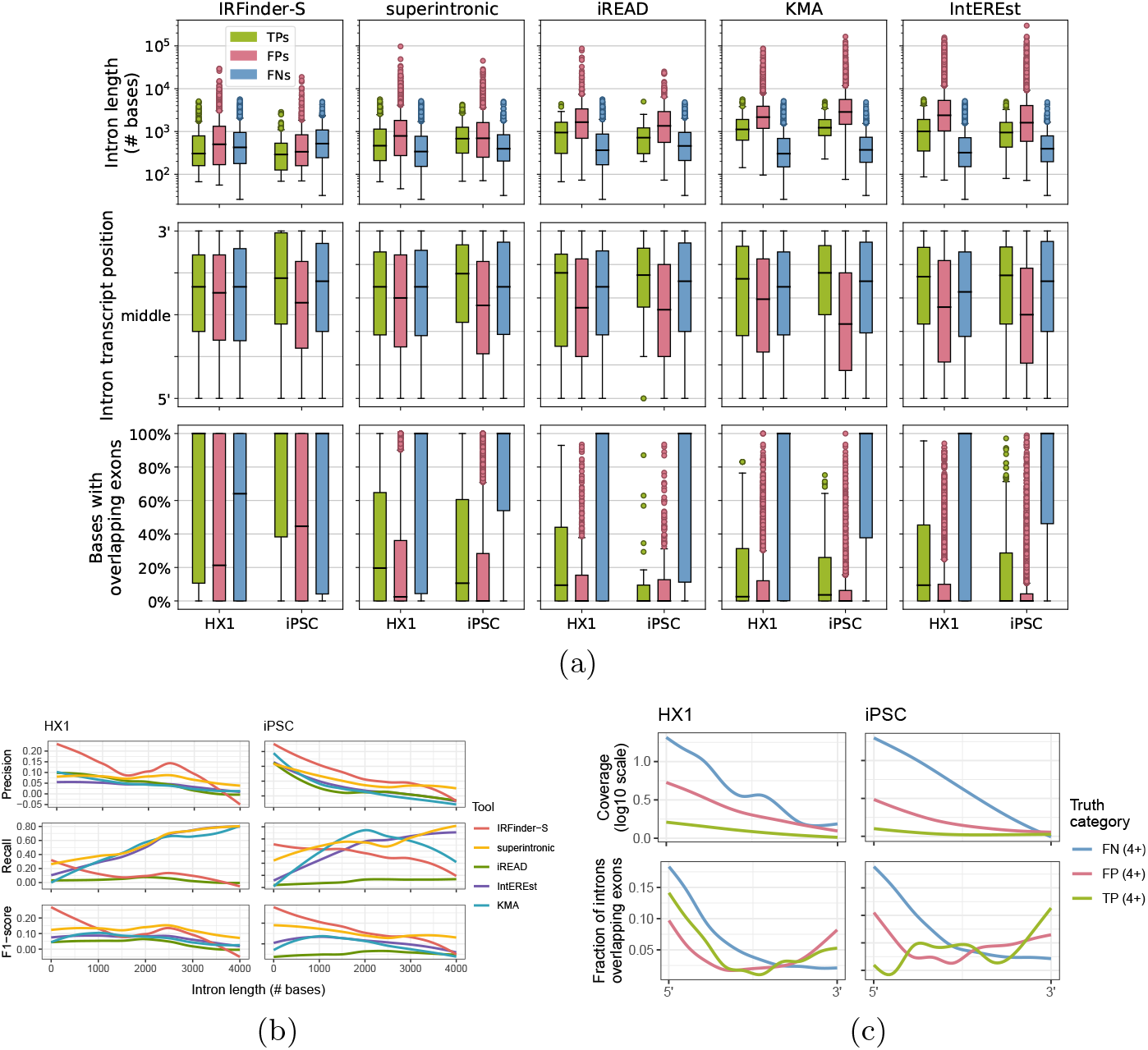
(a) Distributions of properties of TP, FP, and FN RIs across short-read detection tools. Each panel displays the boxplot distributions of intron length (top, log scale), relative position in transcript (middle), and % of intron bases with overlapping annotated exons (bottom) for the output from each of five short-read tools (from left to right: IRFinder-S, superintronic, iREAD, KMA, IntEREst). Y-axes correspond to intron properties as labeled, with each boxplot along the x-axis corresponding to the TP (green, left boxes), FP (pink, middle boxes), and FN (blue, right boxes) calls for HX1 (left) and iPSC (right). (b) Short-read tool performance as a function of intron length. Each panel depicts the LOESS-smoothed precision (top), recall (middle), or F1-score (bottom) in either the HX1 (left) or iPSC (right) sample across overlapping, sliding window intron length ranges (Methods). Smooths are grouped and colored by five short-read tools (red = IRFinder-S, yellow = superintronic, green = iREAD, purple = IntEREst, blue = KMA). (c) Read coverage and exon overlap as a function of position within an intron. LOESS-smoothed short-read data (see Methods) show the median log10-scaled coverage (top row, y-axes) and fractions of introns with overlapping exons (bottom row, y-axes) as a function of position (x-axis, 5’ → 3’ on positive strand) for HX1 (left column) and iPSC (right column). Introns were grouped by truth category membership for at least 4/5 tools (colors, blue = FN, pink = FP, green = TP).

We also probed how much distributional uniformity of mapped read coverage across an intron (coverage “flatness” [39, 41]) and incidence of overlapping exons differed among TPs, FPs, and FNs. Coverage of FPs and to a greater degree FNs was nonuniform, where coverage decreased roughly monotonically from 5’ to 3’ intron ends. Coverage of TPs was comparatively uniform, where coverage was in general substantially lower than for FPs and FNs (Fig. 4c, top two plots). Closer to their 5’ ends, FNs were distinguished by their tendency to overlap exons (Fig. 4c, bottom two plots). Indeed, for superintronic, iREAD, KMA, and IntEREst, the majority of FNs appear to be accounted for by overlapping exons (Fig. 4a). Overlapping exons may thus be a key obstacle to improving recall of many short-read RI detection tools.

### 2.5 Persistent introns or called RIs occur in genes with experimentally validated IR

Finally, we searched the literature and third-party resources for independent evidence of persistent introns appearing in the HX1 and iPSC samples studied here. We examined RI presence in 9 genes (5 in HX1 and 7 in iPSC) that have experimentally validated IR from a variety of cell types and tissues (Table S3) [17, 49–51]. We found that intron retention across these 9 genes varied substantially by sample (no TP introns were observed in both HX1 and iPSC) (Fig. 5). We also found significant variation between the set of RIs in these genes called by different short-read tools, with only a single TP intron in *IGSF8* identified across all tools for iPSC (Fig. S15). Interestingly, the genes *SRSF7* [24, 52] and *AP1G253* [53] appear to be generally enriched for persistent introns, potentially consistent with post-transcriptional splicing [6, 54].

**Fig. 5:**
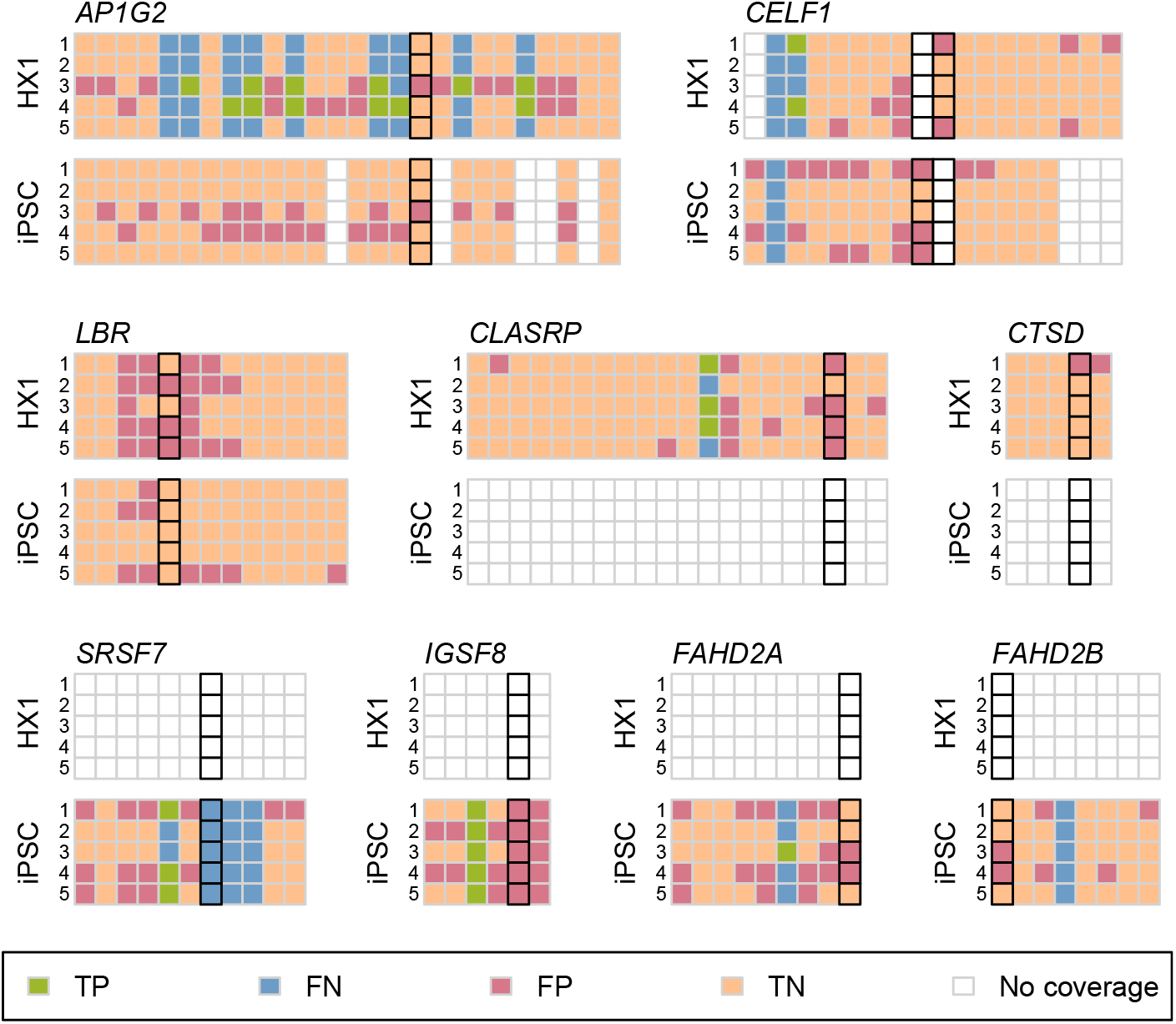
Short-read tool performance across nine genes with experimentally validated RIs. Comparison of short-read tool called RIs with introns detected in long-read data are shown as a pair of matrices for each of nine genes (*AP1G2*, *CELF1*, *LBR*, *CLASRP*, *CTSD*, *SRSF7*, *IGSF8*, *FAHD2A*, and *FAHD2B*). The rows in each matrix correspond to the results from each of five short-read tools (from top to bottom: 1: IntEREst, 2: iREAD, 3:IRFinder-S, 4: superintronic, 5: KMA) applied to either HX1 (top) or iPSC (bottom) data; columns correspond to all introns found across all annotated transcript isoforms of the indicated gene, ordered by left and then right genomic coordinates. Each cell in the matrix depicts the presence or absence of an intron in short-read and/or long-read data as a TP (green), FN (blue), FP (pink), and TN (peach) assessment; white boxes indicate introns found only in transcripts with < 5 assigned long reads. Black outlines indicate the experimentally validated RI(s) in each gene.

## 3 Discussion

This is the first study to evaluate the quality of short-read RI detection using short- and long-read RNA-seq data from the same biological specimen. This study also establishes a novel metric capturing the persistence of an intron in a transcript as it is processed using deep long read RNA-seq, and it is the first to interrogate the potential effects of splicing progression during transcript processing and spurious sources of intronic sequence. We find that short-read tools detect IR with poor recall and even worse precision, calling into question the completeness and validity of a large percentage of putatively retained introns called by commonly used methods. While our results indicate that it may be possible to improve precision slightly by applying expression filters to potential RIs, this appears to come at significant expense to recall.

This work raises fundamental questions regarding how results from short-read RI detection tools should be interpreted. We have taken IR to mean the persistence of an intron in a transcript after processing is complete, in alignment with the biological literature on IR. Short-read RI detection tools are commonly thought to identify such retained introns, with the assumption that poly(A) selection is sufficient to guarantee fully spliced and mature transcripts for sequencing; however, these tools are not inherently designed to distinguish intron retention from contaminating events such as partial transcript processing. This disconnect between how tool developers and tool users employ the same language may be responsible for false assertions in the published literature about which introns are retained. We note, for instance, that the prediction of putative neoepitopes arising from IR [31–35] requires confidence in the detection of stable, persistent IR with a high likelihood of translation and a low likelihood of undergoing NMD, none of which is assured by short-read RI detection tools.

Limitations of this work include the small number of biological specimens with matched short and deep long read RNA-seq available in the public domain, the lack of replicates of short-read RNA-seq data in this setting, and the limited depth of the long read sequencing data. As a result, we were unable to study the patterns of IR across tissue type and other distinguishing sample characteristics. We confined attention to introns that occur in genes with high coverage in both short and long read data, and did not address either confidence in IR as a function of read depth or systematic biases in gene coverage as a function of sequencing platform. While an improvement, our intron persistence metric only partially accounts for admixed splicing patterns from different cell types in a mixed-cell sample such as HX1. Like other RI detection studies [15–17, 25, 31, 34, 43], our approach is explicitly linked to annotation (here, GENCODE v35) and therefore reports IR only relative to annotated transcripts, ignoring potential unannotated transcripts. We also did not explore the entanglement of biological and technical effects in the length of persistent introns: shorter introns are more likely to be retained [43, 55, 56], but the length limit of PacBio Iso-Seq reads of up to 10 kilobases means that any molecules with longer persistent introns were not considered in this study. Furthermore, we calculated length-weighted median expression to harmonize short-read tool outputs to long-read intron ranges (Fig. S16), and this stringent approach may have inflated false negative rates in regions returning high expression magnitudes and variances. Finally, we were only able to evaluate a small subset of the tools available for short read-based RI detection, as many of these tools harbor substantial software implementation and reproducibility challenges.

While there is evidence for cytoplasmic splicing, the phenomenon is rare in many tissues and cell types [9–12]. It may be worth exploring the extent to which sequencing only cytoplasmic RNAs focuses attention on fully processed RNA transcripts in future work.

## 4 Methods

### 4.1 Identification of paired short- and long-read data

Two advanced-search queries were performed on the Sequence Read Archive (SRA) (https://www.ncbi.nlm.nih.gov/sra) on July 13, 2021, and all experiment accession numbers were collected from the query results by downloading the resulting `RunInfo` CSV files. For both searches, the query terms included organism “human,” source “transcriptomic,” strategy “rna seq,” and access “public” with platform varying between the two searches: “pacbio smrt” for the long-read query and “illumina” for the short-read query. The `RunInfo` files were merged and projects with both Illumina and PacBio sequencing performed on the same NCBI biosample were identified. Due to relatively low sequencing depth of PacBio experiments, all projects with fewer than 20 PacBio sequencing runs were eliminated. PacBio experiments conducted on any PacBio platform earlier than RS II were also removed. Two remaining biosamples were chosen as data on which to test RI detection: 1) biosample SAMN07611993, an iPS cell line collected and processed by bioproject PRJNA475610, study SRP098984, with 1 short-read and 27 long-read runs [45], and 2) biosample SAMN04251426 (HX1), a whole blood sample collected and processed by bioproject PRJNA301527, study SRP065930, with 1 short-read and 46 long-read runs [44]. (See the project repository at https://github.com/pdxgx/ri-tests for accession numbers.)

### 4.2 Long-read data collection, initial processing, and alignment

Raw Iso-Seq RS II data were downloaded from the SRA trace site (https://trace.ncbi.nlm.nih.gov/Traces/sra), via the “Original format” links under the “Data access’ tab for each run. These comprised three `.bax.h5` files for both samples, with an additional `.bas.h5` and metadata file for each HX1 run. For both samples, individual runs were processed separately as follows, with differences in handling of the two samples as noted. Subreads were extracted to BAM files from the raw movie files using `bax2bam` (v0.0.8). Circular consensus sequences were extracted using ccs (v3.4.0) with `--minPasses 1` set to `1` and `--minPredictedAccuracy 0.90`. Barcodes were removed from CCS reads and samples were demultiplexed with `lima` (v2.2.0). For HX1, the input barcode FASTA files were generated from the Clontech 5p and NEB Clontech 3p lines from “Example 1” `primer.fasta` (https://github.com/PacificBiosciences/IsoSeq/blob/master/isoseq-deduplication.md). For iPSC, forward and reverse barcode fasta files were downloaded from the study’s GitHub page (https://github.com/EichlerLab/isoseqpipeline/tree/master/data) and merged into a single FASTA file per the `lima` input requirements. Since lima generates an output file for each 5’-3’ primer set, these were merged using `samtools merge` (`samtools` and `htslib` v1.9). Demultiplexed reads were refined and poly(A) tails removed using `isoseq3 refine` (`isoseq` v3.4.0) to generate full-length non-concatemer (FLNC) reads. FLNC reads were extracted to FASTQ files using `bedtools bamtofastq` (`bedtools` v2.30.0), and aligned to GRCh38 with `minimap2` (v2.20-r1061) using the setting `-ax splice:hq`. Sequence download and processing scripts are available at https://github.com/pdxgx/ri-tests. After processing, the 46 HX1 Iso-Seq runs yielded 945,180 aligned long reads covering 32,837 transcripts of 11,813 genes for HX1, with 13,560 of these transcripts covered by at least 5 long reads and 4409 unique 5+ read transcripts showing evidence of possible intron retention. In the iPSC sample, we obtained 839,558 aligned long reads covering 31,546 transcripts of 11,992 genes. 12,676 of these transcripts were covered by at least 5 long reads, with 3137 unique 5+ read transcripts showing evidence of possible intron retention.

### 4.3 Assignment of long reads to transcripts

The long-read alignment files were parsed as follows. GENCODE v.3557 annotated transcripts’ introns, strand, and start/end positions were extracted from the GENODE v35 GTF file. Then for each aligned long read, spliced-out introns, strand and start/end positions were extracted using `pysam` (v0.16.0.1, using `samtools` v1.10) [57, 58]. A set of possible annotated transcripts was generated, comprising transcripts for which the read’s set of introns exactly matched the annotated transcripts’ introns sets (“all introns”), or if no such transcripts were found, transcripts for which the read’s introns were a subset of the transcripts’ intron sets (“skipped splicing”). Then the best transcript match was chosen from the shortlist of potential matches as the transcript whose length most closely matched the read length. Some reads did not cover the full lengths of their best-matched transcripts, defined by the read alignment start and end position encompassing all introns in the annotated transcript (“full length”); in the case where not all intron coordinates were covered, these were labeled “partial” reads.

### 4.4 Intron persistence calculation

Intron persistence was calculated only for every transcript that was assigned as the best match for at least 5 reads. We calculated persistence for each intron within these transcripts as the information density of the intron *d_i_* (i.e., the proportion of reads assigned to the transcript that cover intron *i*) multiplied by the mean of the product of three terms across all long reads assigned to that isoform:

1. The **retention**, or presence, *R_r,i_* of a given intron *i* is 1 if the read wholly contains *i* or 0 if it is absent/spliced out as annotated in read *r*.
2. The **spliced fraction** (*SF_r,i_*) for a given intron *i* and read *r* is defined as

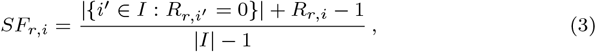

where *I* is the set of introns spanned by *r* and *R_r,i_* is defined above. This fraction of spliced introns in a read, with the target intron excluded, represents the splicing progression of the read. A mature RNA molecule should tend to have fewer unspliced introns present than an RNA from the same transcript at an earlier point in splicing progression.
3. The scaled **Hamming similarity** (*H_r,i_*) for a given read *r* and intron *i* is defined as the average number of spliced or unspliced introns that match between the target read and other reads assigned to the transcript that have intron *i* spliced the same as in read *r*, scaled to the number of introns in the isoform:

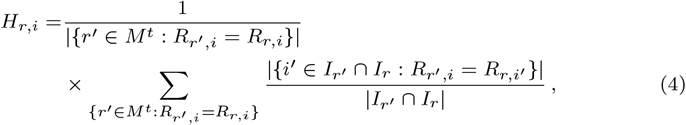

where *I_r_* is the set of introns spanned by *r*, *I_r′_* ∩ *I_r_* is the set of introns covered by both *r* and *r*′, *M ^t^* is the set of reads assigned as best matches to the same transcript as *r* and span the target intron *i*, and *R_r,i′_* is as defined above. Any partial reads that are assigned to the transcript as a best match but do not span the target intron are not included in this calculation, and the scaled Hamming similarity between two reads is only calculated for introns covered by both reads. This term accounts for the stochasticity of splicing initiation and progression, since a collection of reads would be more likely to have a dissimilar pattern of unspliced introns if the splicing process remained incomplete.

Persistence *P_i,t_* was calculated for each intron *i* in a given transcript isoform *t* as information density of the intron *d_i_* times the mean of the product of the three terms above per Equation 1. Since short reads are not assignable to specific transcripts or isoforms, and certain introns fully or partially recur across multiple transcripts, we set the **intron persistence** *P_i_* for a given intron i as the maximum *P_i,t_* found for that intron across all transcripts in which it occurs per Equation 2.

### 4.5 Alignment and BAM generation for short-read data

FASTQs were generated using either Illumina’s NextSeq 500 (iPSC, run id: SRR6026510) or HiSeq 2000 (HX1, run id: SRR2911306), and files were obtained from the SRA using the `fastq-dump` command from the SRA Toolkit (v2.10.8). A `STAR` (v2.7.6a) [59] index was generated based on the GRCh38 primary assembly genome FASTA (ftp://ftp.ebi.ac.uk/pub/databases/gencode/Gencode_human/release_35/GRCh35.primary_assembly.genome.fa.gz) and GTF (ftp://ftp.ebi.ac.uk/pub/databases/gencode/Gencode_human/release_35/gencode.v35.primary_assembly.annotation.gtf.gz) files from GENCODE v35 [60]. Reads were aligned with STAR to this index using the `--outSAMstrandField intronMotif` option. Primary alignments were retained for reads mapping to multiple genome regions. SAM files output by `STAR` were converted to both sorted and unsorted BAM files using `samtools sort` and `view` (`samtools` v1.3.1), respectively.

Additionally, for use with KMA [37], `bowtie2` (v2.3.4.3) [61] alignments were performed. Alignment statistics may be found in the project repository (https://github.com/pdxgx/ri-tests) and are summarized in Fig. S17. A FASTA file with intron sequences was generated based on the GRCh38 primary assembly genome FASTA and GTF files from GENCODE version 35 using the `generate_introns.py` script from the KMA package setting 0 bp for the `extension` flag. These intron sequences were combined with the GRCh38 transcript sequence FASTA file from GENCODE version 35 (ftp://ftp.ebi.ac.uk/pub/databases/gencode/Gencode_human/release_35/gencode.v35.transcripts.fa.gz), and this combined FASTA was used to create a Bowtie 2 index. Reads were aligned to this index using `bowtie2` according to specifications from KMA [62]. To quantify expression from the Bowtie 2 alignments, eXpress (v1.5.1) [63, 64] was used.

### 4.6 Selection of target gene subset

Due to variable short- and long-read coverage across the genome, we selected a subset of genes to use for our test dataset to ensure adequate sequencing coverage for RI detection on both platforms. For the short-read data, we chose a coverage cutoff based on the requirements of the short-read RI detection tools used. The two tools with clear coverage requirements are iREAD, which requires coverage of 20 reads across an intron for RI detection, and superintronic, which requires 3 reads per region. Since these are short-reads (126 bases for iPSC and 90 for HX1) required over potentially long intronic regions, we chose a median gene-wide coverage (including both intronic and exonic regions) of 2 reads per base, ensuring either consistent coverage across the gene or high coverage in some areas. For the PacBio data, we selected 5 long reads per gene, and a further filter of at least 5 reads aligned to a single transcript of the gene, as giving enough information for comparing splicing progression and splicing patterns between reads. The target gene sets, 4,639 genes for iPSC and 4,369 for HX1, were chosen from the aligned data, naive to potential RI detection, and then for both short- and long-read data, the gene subset was applied as a filter after running metric calculations or RI detection by short read tools. Within these genes, only transcripts with at least 5 long reads were studied.

### 4.7 Intron feature annotation

For the set of target genes, transcripts with at least 5 long reads were selected for analysis. Features of each intron in these transcripts including intron lengths, splice motif sequences, relative transcript position, spliceosome category, and transcript feature overlap properties were extracted as follows. Length was calculated as the difference between the right and left genomic coordinates of the intron ends. Relative position within the transcript is an intron-count normalized fraction where 0 represents the transcript’s 5’ end and 1 represents the 3’ end. Splice motifs were assigned to each intron by querying the GRCh38 reference genome with `samtools faidx` (`samtools` v1.10) for the two coordinate positions at each end of the intron, and assigned to one of three canonical motif sequences (GT-AG, GC-AG, and AT-AC, and their reverse complements for − strand genes) or labeled as “other” for noncanonical motifs. Three feature overlap properties were studied: the total number of exons from other transcripts with any overlap of the intron region; the percent of intron bases with at least one overlapping exon from another transcript; and the maximum number of exons overlapping a single base in the intron. These were calculated by extracting all exon coordinates from the GENCODE v35 annotation file, and using an interval tree to query each intron base position against the set of annotated exon coordinates. Spliceosome category was determined from recent U2 and U12 intron annotations [7]. BED files of U2 and U12 introns for GRCh38 were downloaded from the Intron Annotation and Orthology Database (https://introndb.lerner.ccf.org/) on 1/25/22. Introns were labeled “U2” or “U12” if they only overlapped ranges from one of either spliceosome category, and remaining introns were labeled “other.”

### 4.8 Selection of short-read RI detection algorithms and identification of likely RIs

We successfully downloaded and ran five IR-specific detection tools for short-read data on our remote server using the CentOS v7 operating system. To run superintronic, KMA, IntEREst, and iREAD, we used conda virtual environments (see https://github.com/pdxgx/ri-tests). We ran IRFinder-S from a fully self-contained Dropbox image per the tool’s instructions (see below). IntEREst and superintronic are provided as R libraries which we ran from interactive R sessions, while iREAD, IRFinder-S, and KMA were run from command line, and a separate R package was used for RI detection for KMA. Outputs from all tools were read into R and harmonized to a single set of intron ranges after applying minimum coverage filters based on both short-read and long-read data. After running tools according to their provided documentation, we consulted literature and documentation on a tool-by-tool basis to devise starting filter criteria based on expression magnitude and other properties. We used these starting criteria to find the subset of most likely RIs, then we modified filter criteria to ensure filtered intron quantities were roughly one order of magnitude lower than unfiltered introns in both iPSC and HX1.

### 4.9 IR quantification with IntEREst

To run IntEREst (v1.6.2) [38], the `referencePrepare` function from the package was used to generate a reference from the GENCODE v35 primary assembly GTF file [60]. This reference was used along with the sorted STAR BAM alignment from each sample to detect intron retention with the `interest` function, considering all reads and not just those that map to junctions. We used the `interest` function with the `IntRet` setting, which takes into account both intron-spanning and intron-exon junction reads and returns expression as a normalized FPKM. The filter FPKM ≥ 3, recommended for iREAD, left > 90% of introns in both samples, so we increased the minimum filter to FPKM ≥ 45, and this retained 5038/32544 ≈ 15% of introns in HX1 and 6832/21820 ≈ 31% of introns in iPSC (Fig. S10).

### 4.10 IR quantification with keep me around (KMA)

To run KMA [37], we used devtools to install a patched version of the software which resolves a bug unaddressed by the authors, available at https://github.com/adamtongji/kma. The `read_express` function was used to load expression quantification data output from eXpress, and the `newIntronRetention` function was used to detect intron retention. Returned intron expression was scaled as transcripts per million (TPM). We noted the recommended filters of unique counts ≥ 3 and TPM ≥ 1 left just 7.2% of introns in iPSC versus 19% in HX1, so we used a less stringent filter of unique counts ≥ 10 for both samples, which left 6437/14155 ≈ 45% of introns in iPSC and 5089/20484 ≈ 25% of introns in HX1 (Fig. S10).

### 4.11 IR quantification with iREAD

To run iREAD (v0.8.5) [39], a custom intron BED file was made from the GENCODE v35 primary assembly GTF file using GTFtools (v0.6.9) [65]. The total number of mapped reads in each sorted STAR BAM alignment was determined using samtools, and used as input to the iREAD python script to detect intron retention. Intron expression was returned scaled as FPKM. To identify the most likely RIs, we applied previously published filter recommendations for entropy score (≥ 0.9) and junction reads (≥ 1). Since there were relatively few introns remaining after applying published filters to the iPSC short-read data (313/19316 ≈ 1.6% vs. 583/7748 ≈ 7.5% in HX1), we applied lower filters for FPKM (≥ 1 vs. ≥ 3) and read fragments (≥ 10 vs. ≥ 20) (Fig. S10).

### 4.12 IR quantification with superintronic

To run superintronic (v0.99.4) [40], intronic and exonic regions were gathered from the GENCODE v35 primary assembly GTF file [60] using the `collect_parts` function. The `compute_coverage` function was used to compute coverage scores for each sample from sorted STAR BAM alignments, and the `join_parts` function was used to convert these scores to per-feature coverage scores. Intron expression was returned as log_2_-scaled coverage, and we identified retained intron ranges as those overlapping long read-normalized ranges with LWM ≥ 3, per the expressed introns filter described in [40] (Fig. S10).

### 4.13 IR quantification with IRFinder-S

We ran IRFinder-S v2.0-beta using the Docker image obtained from https://github.com/RitchieLabIGH/IRFinder. We prepared the IRFinder reference files using the GENCODE v35 genome sequence reference and intron annotations [60]. Our analyses focused on the coverage and IRratio metrics, and the intron expression profile flags returned under warnings. Intron expression was returned as an IRratio, which is similar to PSI, and we identified likely retained introns as having IRratio ≥ 0.5 without any flags per the methods in [41] (Fig. S10).

### 4.14 Harmonization of intron retention metrics across algorithms and runs

Prior to analysis, we harmonized algorithm outputs on intron ranges returned by analysis of available long read runs. We harmonized intron expressions from short read RI detection tools to intron ranges remaining after long reads were uniquely mapped to transcript isoforms. For each short-read RI detection tool, we calculated the region median intron expression value after weighting values on overlapping range lengths (a.k.a. length-weighted medians [LWM]). Calculation of LWMs is shown for an example intron in Fig. S16. Interrater agreement among the output from different short-read algorithms was assessed by Fleiss’ kappa [66] using the R package `irr` v0.84.1.67 [67].

### 4.15 Calculation of performances by intron length bins

We calculated called RI performance metrics across five short-read tools for a series of overlapping intron length bins. In total, 41 bins were calculated for each sample by sliding 300 bp-wide windows from 0 to 4300 bp lengths at 100 bp intervals. Plots were generated by computing LOESS smooths of the binned performance results.

### 4.16 Calculation of normalized binned coverages

We evaluated binned intron characteristics across intron truth metric categories for each sample. We assigned introns to truth categories if they were recurrent in that category for ≥ 4 of 5 short-read tools (e.g., an intron that was recurrent TP for four tools in iPSC, etc.). We then calculated the log10 median short-read coverage for 1,000 evenly spaced bins per intron for each truth category. We further calculated percent of introns overlapping an exon for each bin by using annotations from the GENCODE v35 GTF. Plots were generated by computing the LOESS smooths of the binned results.

### 4.17 Comparison of detected RIs with circular RNA and validated RIs

We downloaded a database of human circular RNAs from circbase [46] (http://www.circbase.org/download/hsa_hg19_circRNA.txt), most recently updated in 2017. We extracted all cRNAs labeled with the “intronic” flag in the annotation column and performed a liftover of genomic coordinates for these cRNAs from hg19 to GRCh38 using the UCSC Genome Browser `liftover` tool (https://genome.ucsc.edu/cgi-bin/hgLiftOver). For each sample, we determined the percent of introns overlapping at least one cRNA for the 4+ consensus truth metric groups TP, FP, and FN (e.g., intron was TP in ≥ 4 tools, etc.)

In order to test introns in this study against experimentally validated RIs, we identified wet-lab studies in the literature that had first predicted, and then validated intron retention. We identified 4 such studies [17, 49–51] that validated a total of 9 RIs in our sets of target genes as defined above (5 and 7 in HX1 and iPSC respectively) (Table S3). (The above four plus an additional ten studies [9, 30, 68–75] experimentally validated RIs in an additional 6 and 9 genes that were found in our target gene sets for in HX1 and iPSC respectively, but without evidence of IR in our samples, and 41 and 36 genes, respectively, that did not pass our sample coverage thresholds for inclusion in this study.) The validated intron coordinates (Table S3) were extracted either from the published intron number [17, 49, 50], assuming a count from the gene’s 5’ to 3’ end, or via BLAT queries of the target sequence [51]. In each sample and tool, we determined the truth status (TP, FP, TN, or FN) of all introns of all transcripts in the target gene for transcripts with ≥ 5 long reads. Adequate intron expression information was available in both samples for the genes *LBR*, *CELF1*, *AB1G2*, but only one sample each for remaining genes.

## Supporting information

Supplementary Performance Data

## Acknowledgments

We thank Kasper Hansen, Jeremy Goecks, and Joe Gray for their helpful feedback as this work was being prepared.

## f Disclaimer

The contents do not represent the views of the U.S. Department of Veterans Affairs or the United States Government.

## Supplementary information

### Supplementary figures

**Fig. S1:**
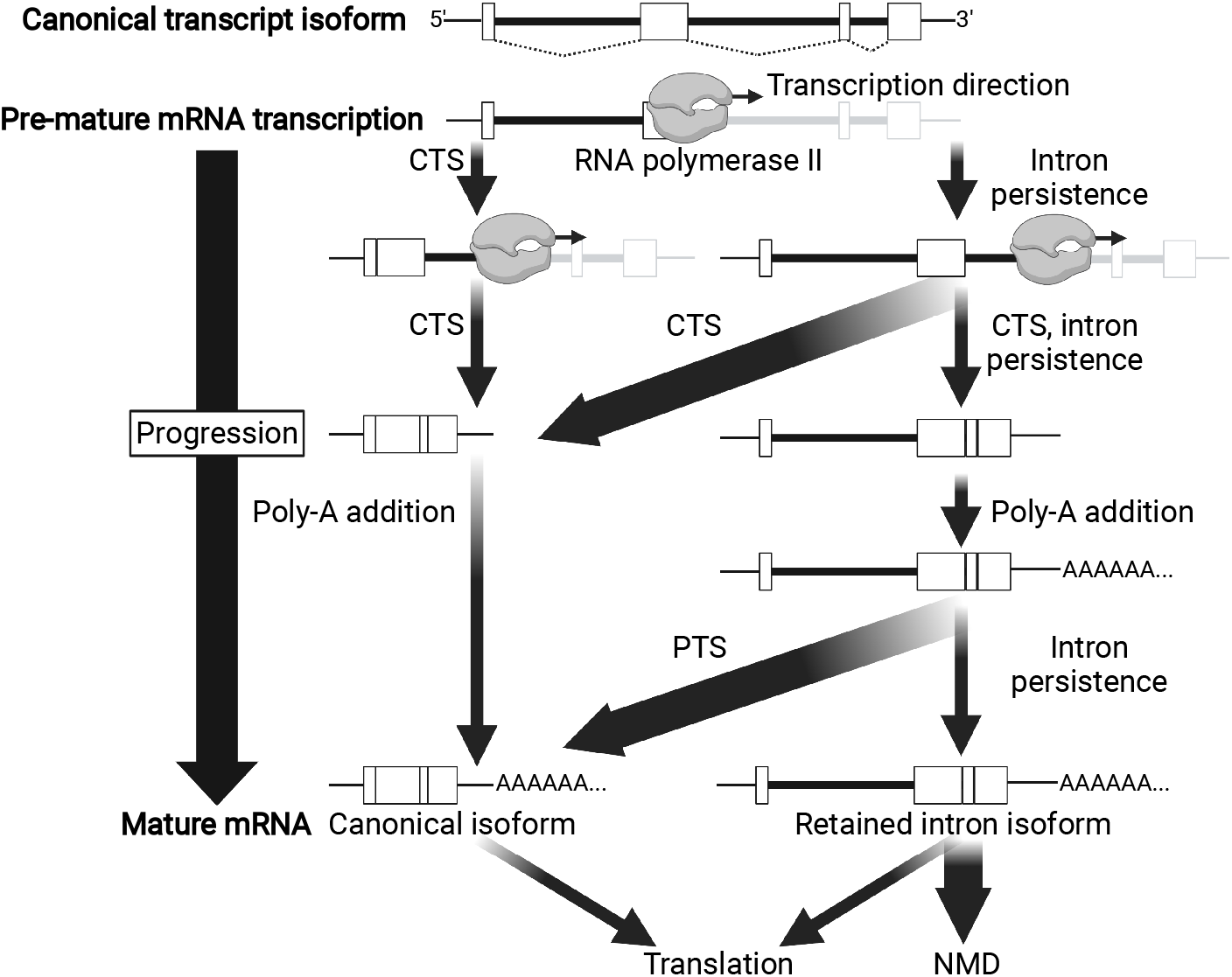
Progression of transcript diagram, created with BioRender.com. Diagram depicts successive steps in transcript processing which progress from top to bottom. At the top is shown the presumed canonical transcript isoform with its expected splice pattern, followed by pre-mRNA processing steps, which branch between transcription by RNA polymerase II, co-transcriptional splicing (CTS), intron persistence, and poly(A) addition. At the bottom are the possible mature mRNA endpoints, including results from post-transcriptional mRNA splicing (PTS) and processing, which include translation and nonsense-mediated decay (NMD). Arrows are labeled with the events they represent, where arrow width sizes indicate their expected event frequencies.

**Fig. S2:**
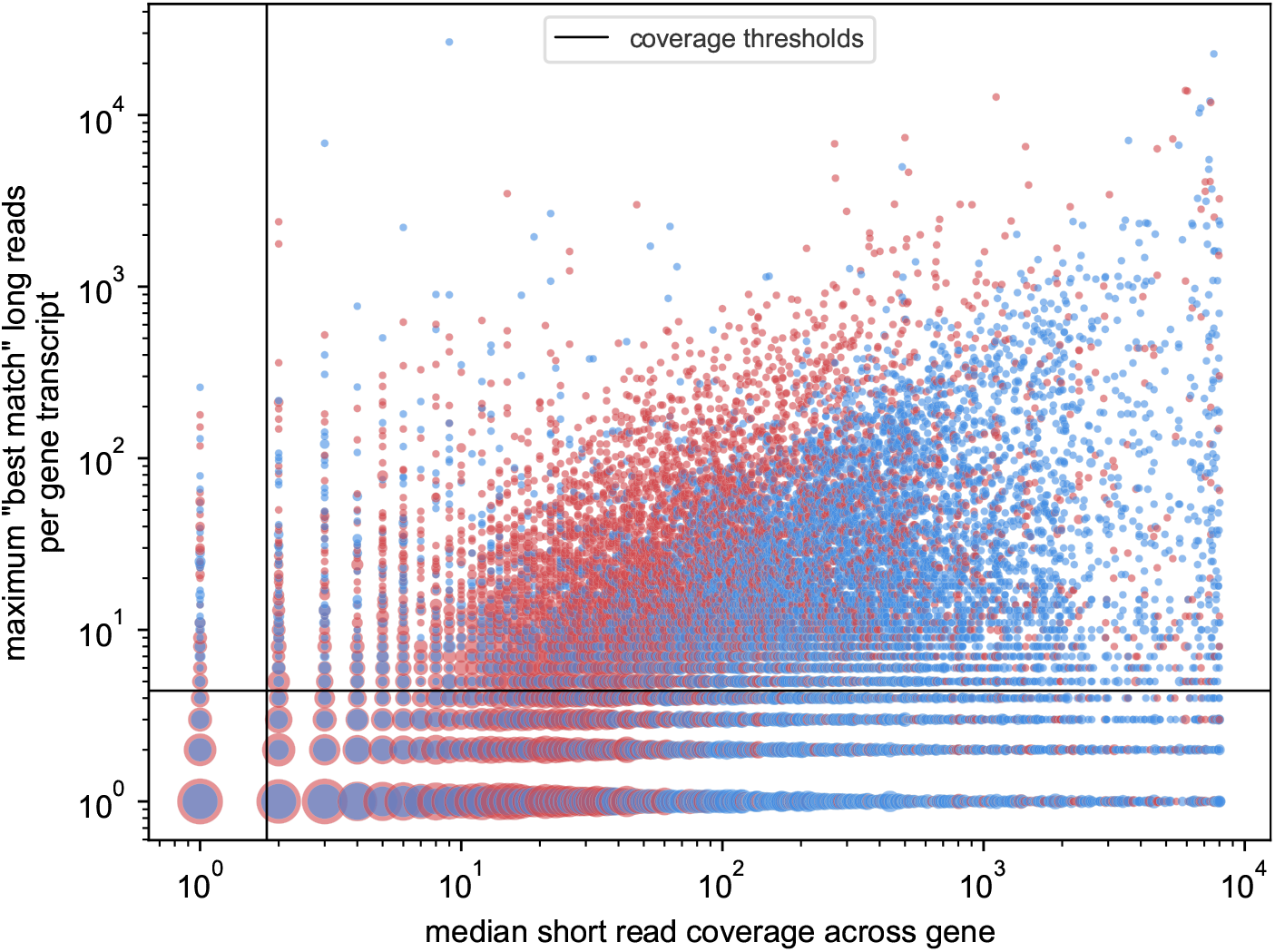
Short- and long-read coverage of genes by sample. The maximum number of long reads assigned to one transcript of each gene (y-axis) vs. the median short-read coverage per base across the entire gene (x-axis) for HX1 (red) and iPSC (blue) samples, in log scale. The vertical line represents the minimum median short-read coverage (2) and the horizontal line represents the minimum total long-read coverage per transcript (5) required for a gene to be included in our analysis; genes considered are in the upper right region of the plot.

**Fig. S3:**
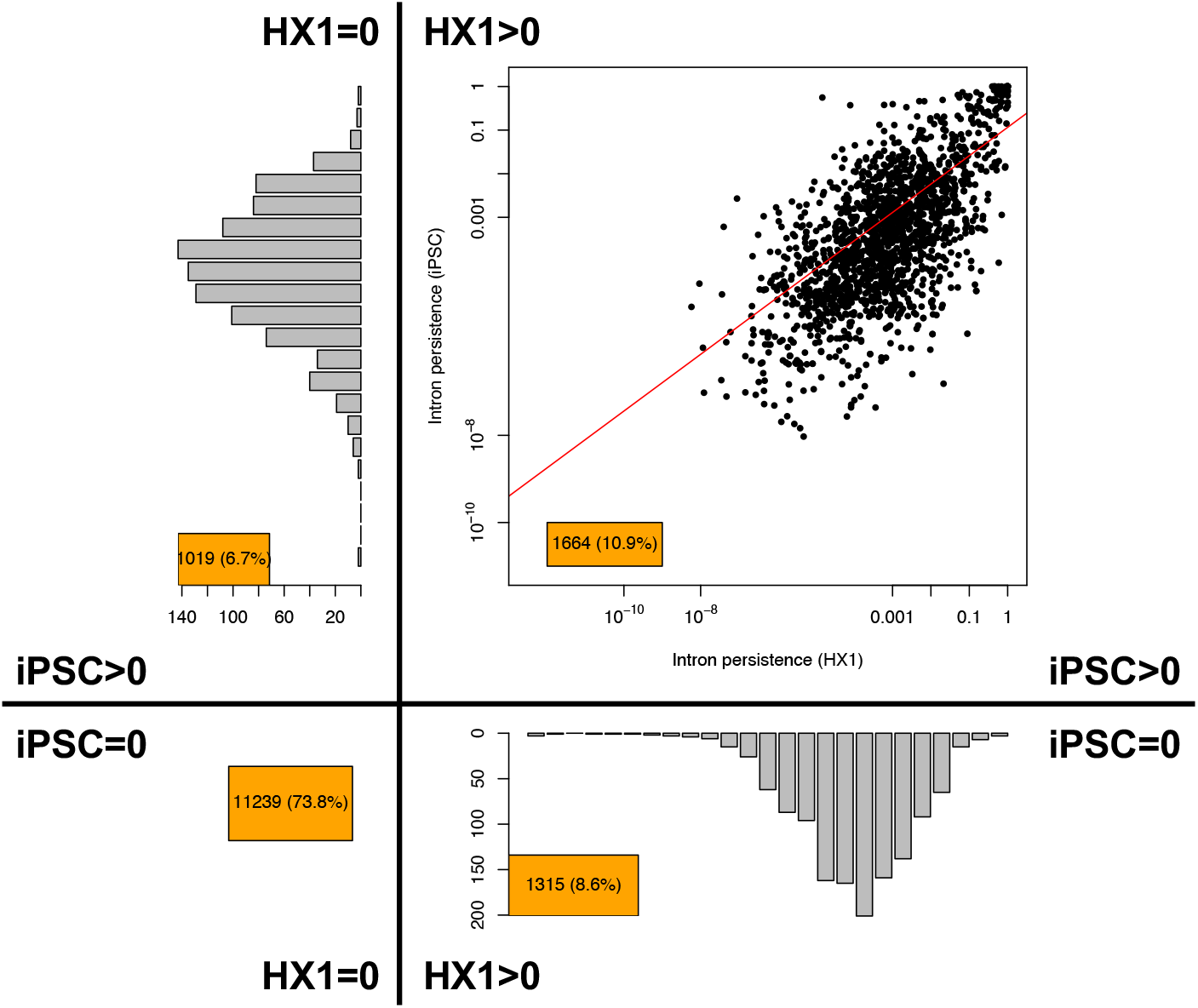
Distribution of intron persistence values for introns in HX1 and iPSC samples. For introns included in both sample studies, bottom left quadrant represents introns with no persistence across both samples (73.8%), upper left represents introns with persistence in iPSC but not HX1 (6.7%), bottom right represents introns with persistence in HX1 but not iPSC (8.6%), and upper right is a scatterplot of persistences in iPSC (y-axis) vs. HX1 (x-axis) for introns with persistence in both (10.9%).

**Fig. S4:**
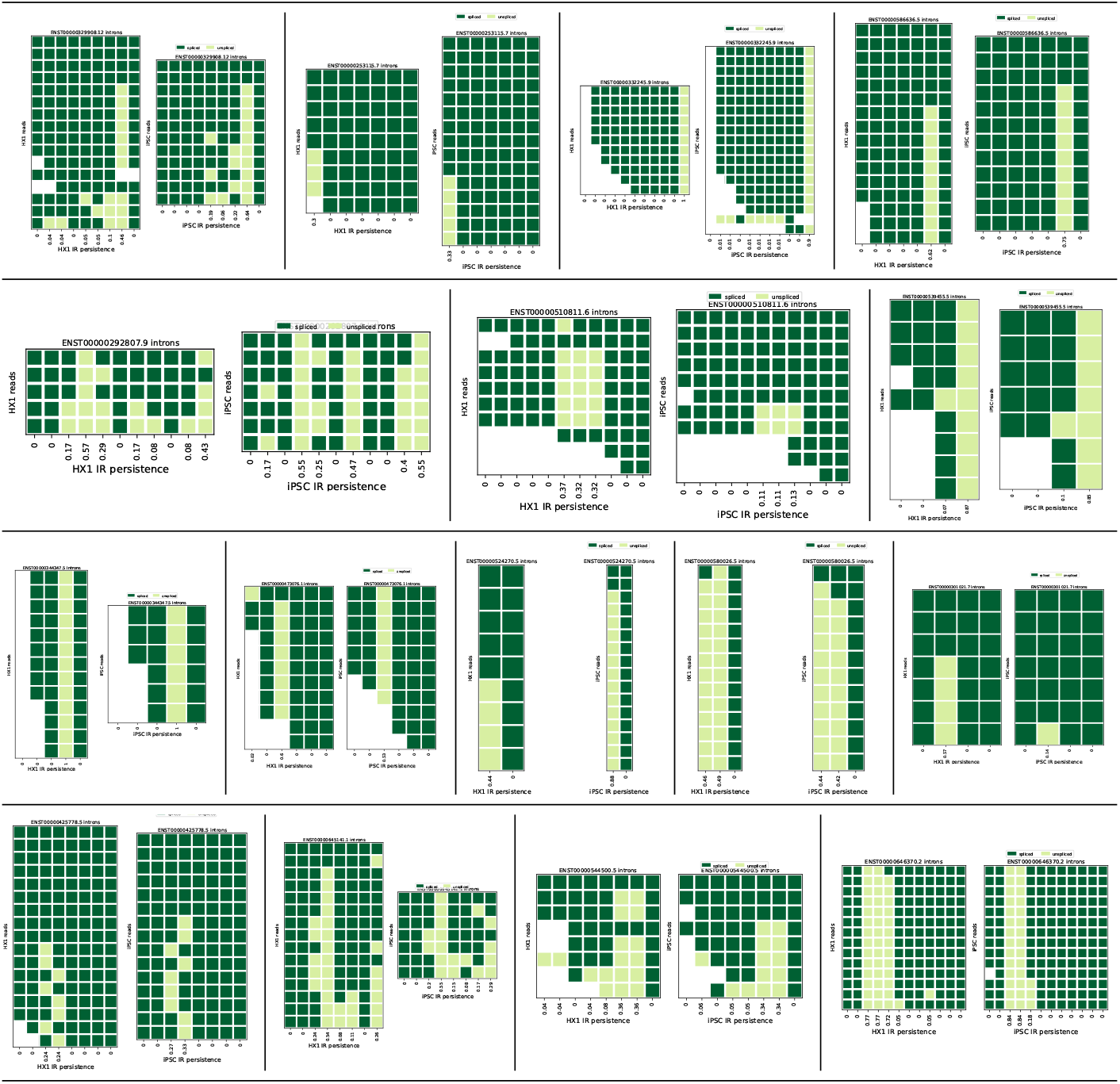
Splicing similarity between samples. Heatmaps of splicing patterns for a selection of matched transcripts between HX1 and iPSC. Each subplot shows data for one transcript in HX1 (left) and iPSC (right) with rows representing transcript-matched long reads and columns representing transcript introns in 5’ → 3’ order. Transcripts were selected from a subset with 5–20 matched long reads in each sample and 5–20 introns. Dark green indicates a spliced out intron in a given read, light green indicates a retained intron, and white indicates no coverage of the intron in the read.

**Fig. S5:**
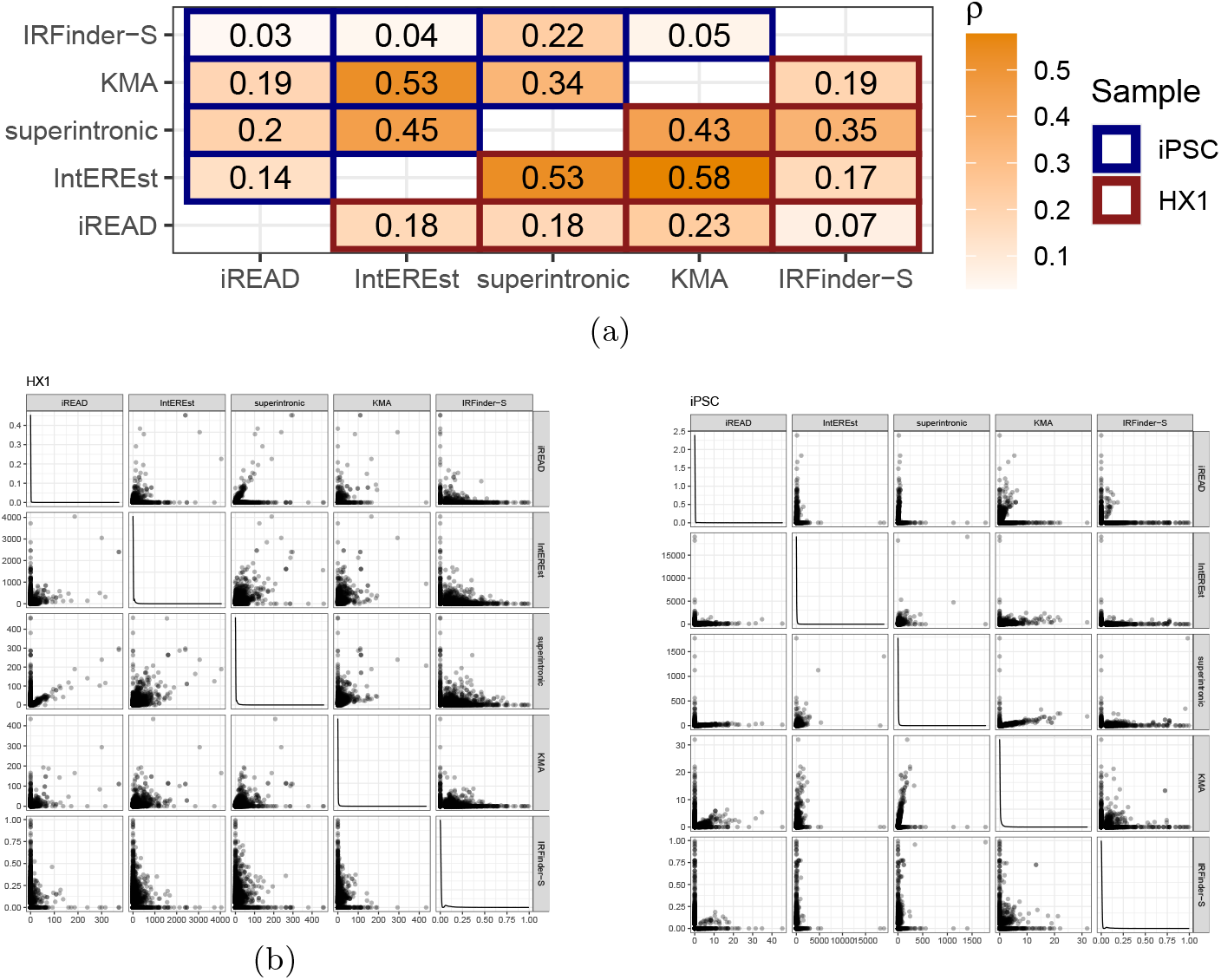
(a) Pairwise correlations among the intron expression values output by five short-read tools. Each element in this heatmap depicts the correlation in intron expression values (Spearman’s test) between the indicated pair of short-read tools, as labeled along the x- and y-axes. Cell text indicates Spearman *ρ* coefficient, with corresponding color value obtained by the color gradient scale shown (from white to orange). Cell outline color indicates the sample for which inter-tool correlation was assessed (iPSC [top left] and HX1 [bottom right] are outlined in blue and red, respectively). (b) Intron expression scatter plots between all short-read IR-detection tool pairs (lower and upper triangles of plot grid) and density plots for each of the five individual tools (diagonal plot grid) for HX1. (c) Intron expression scatter plots between all short-read IR-detection tool pairs (lower and upper triangles of plot grid) and density plots for each of the five individual tools (diagonal plot grid) for iPSC.

**Fig. S6:**
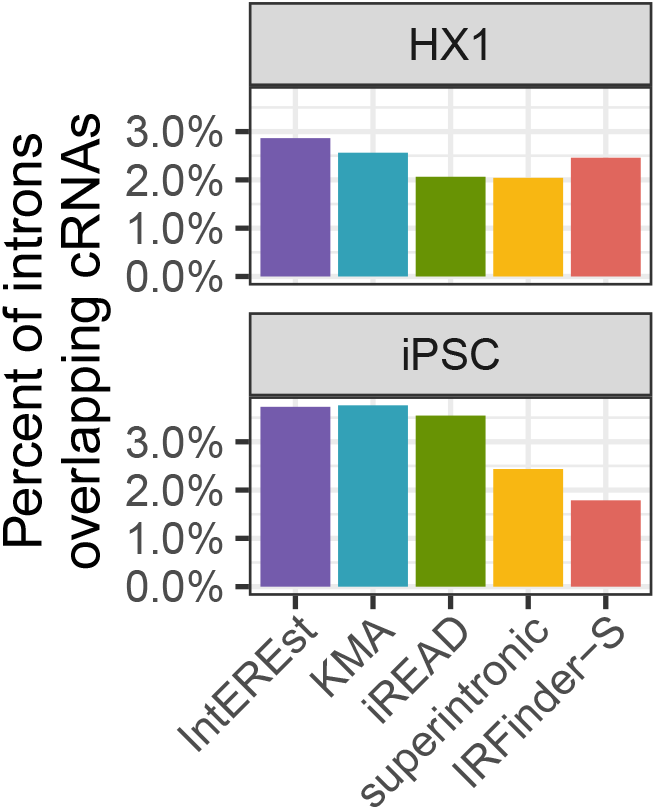
cRNA overlap among called RIs. Barplots quantify the percent of introns overlapping cRNAs (y-axes) across RIs called from five short-read tools (x-axes, bar color fills, red = IRFinder-S, yellow = superintronic, green = iREAD, purple = IntEREst, blue = KMA).

**Fig. S7:**
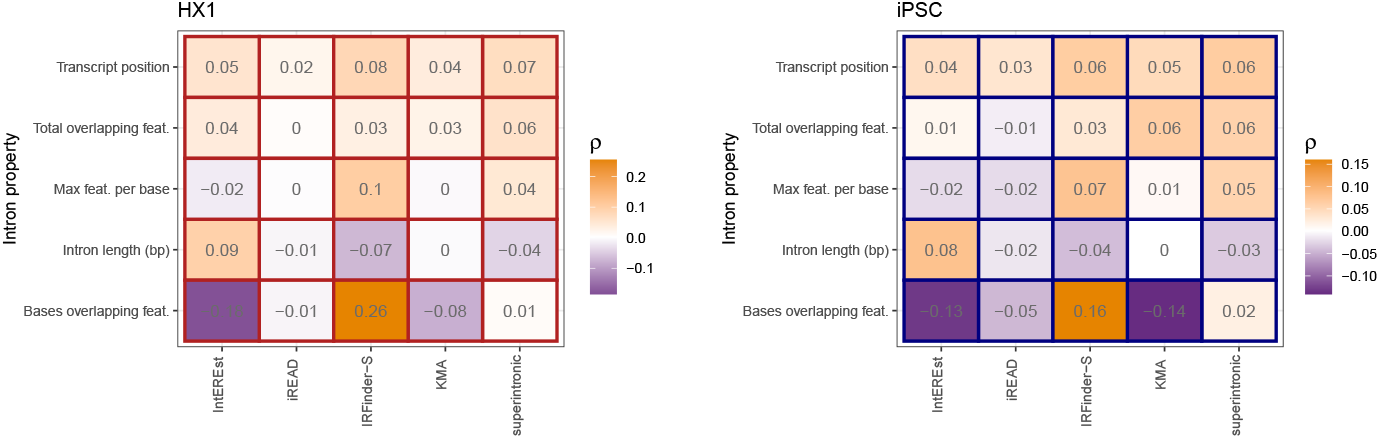
Correlation of intron expression and continuous properties. Heatmap color fills and text show the Spearman *ρ* (purple = negative, white = near zero, orange = positive) between intron expression at five short-read tools (x-axes/columns) and five continuous intron properties (y-axes/rows), for samples HX1 (left heatmap) and iPSC (right heatmap).

**Fig. S8:**
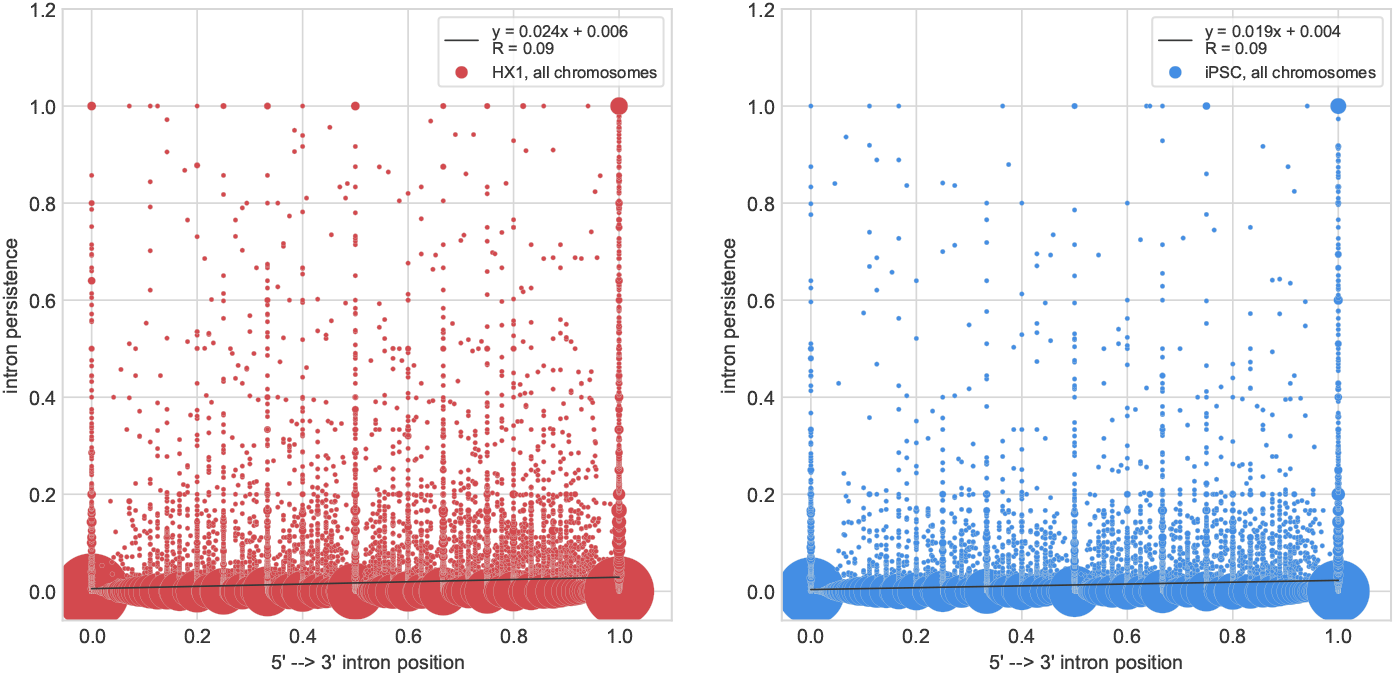
Association of persistence with transcript position. Scatterplots of intron persistence vs. position within a transcript for HX1 (left, red), iPSC (right, blue). Each point represents one or more introns, with point size representing the number of points at each coordinate. Intron position is an intron-count normalized fraction where 0 represents the transcript’s 5’ end and 1 represents the 3’ end. Plotted lines show the linear fit with equations show in the inset legends.

**Fig. S9:**
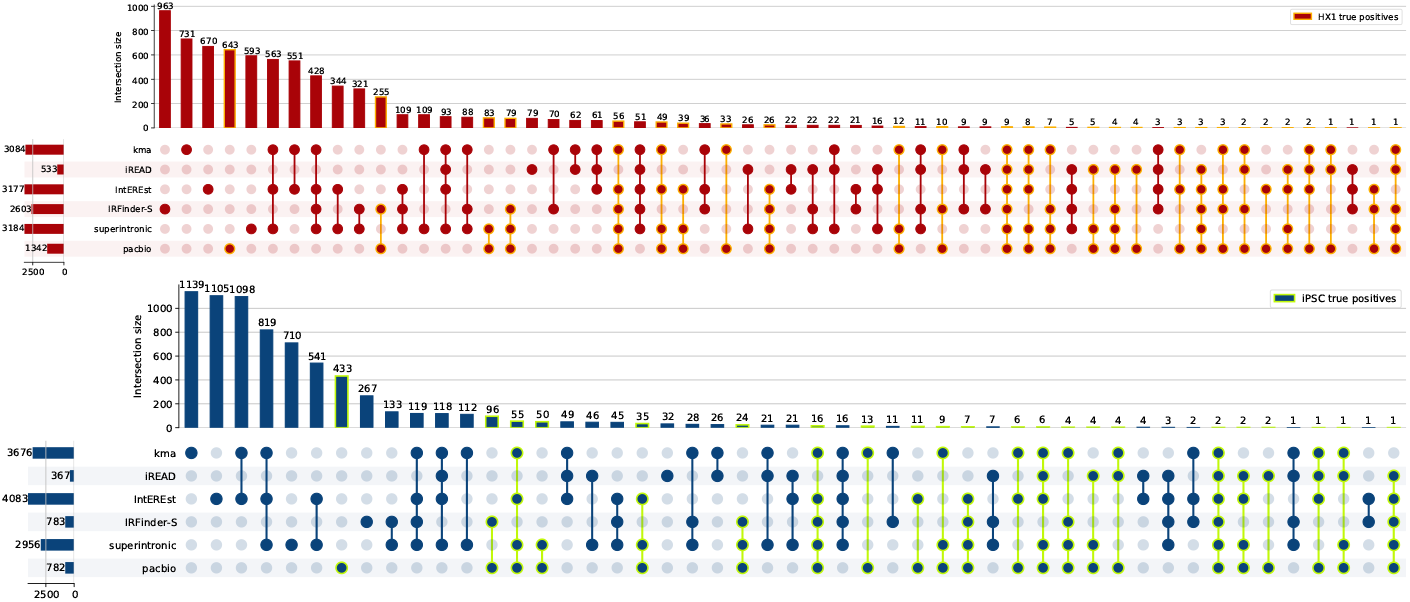
Set overlaps of persistent introns and called RIs. Upset plots showing overlaps of sets of short-read called RIs and long read persistent introns for iPSC (above, blue) and HX1 (below, red). Sets of true positive persistent introns are highlighted in green for iPSC (above) and orange for HX1 (below).

**Fig. S10:**
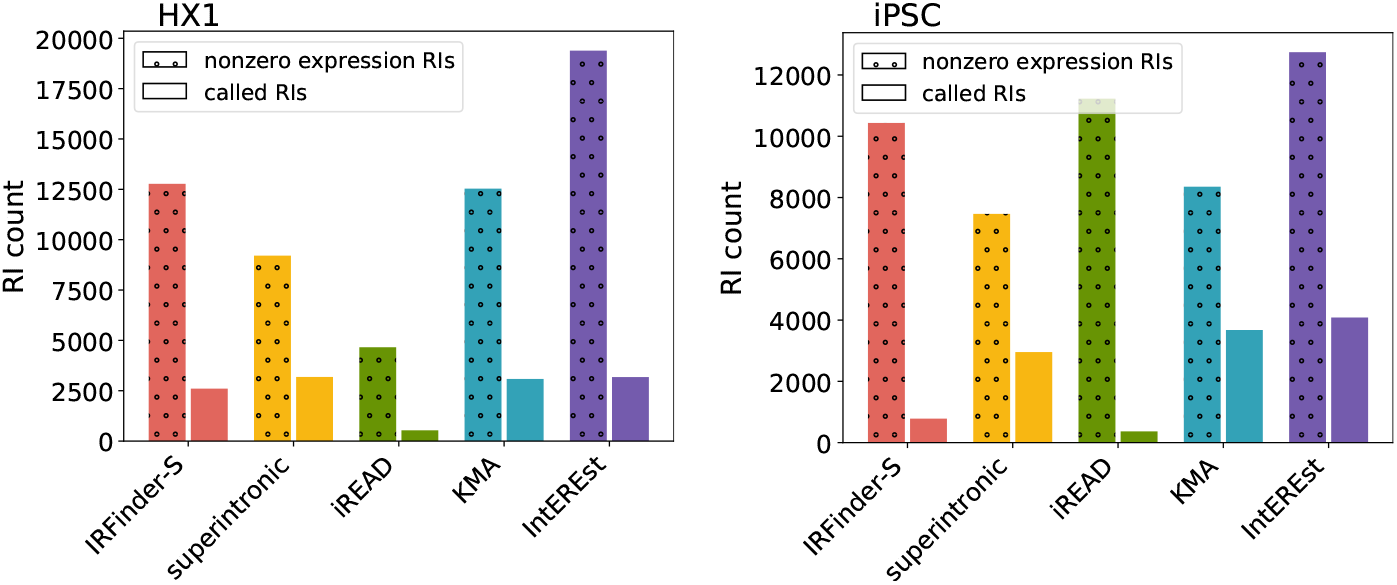
Potential vs. called RI sets. For HX1 (left) and iPSC (right), counts of all potential (calculated nonzero expression) RIs (dotted hatch, left for each tool) and called (filtered) RIs (no hatch, right for each tool) for each SR detection tool (red = IRFinder-S, yellow = superintronic, green = iREAD, purple = IntEREst, blue = KMA).

**Fig. S11:**
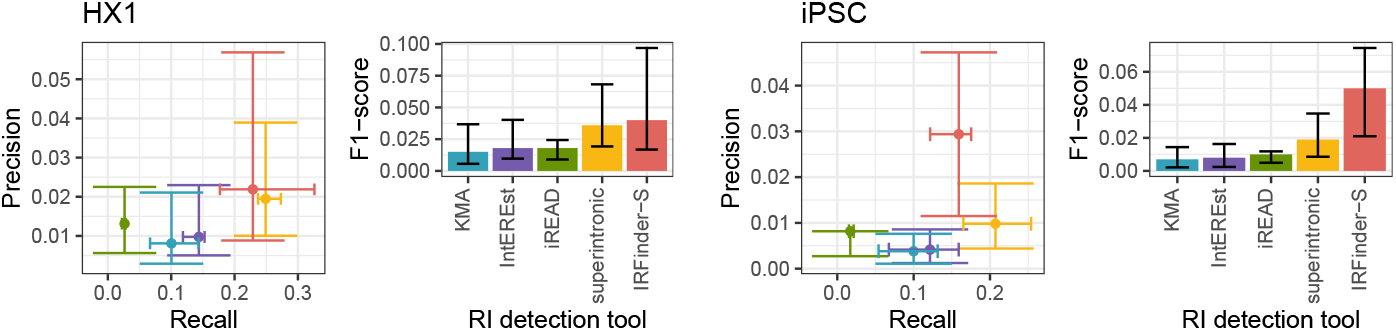
Performance summaries across persistence cutoffs. Scatter plot y-axes show precision and x-axes show recall, and barplot y-axes show F1-scores, for samples HX1 (left plots) and iPSC (right plots). Colors indicate short-read RI detection tools (red = IRFinder-S, yellow = superintronic, green = iREAD, purple = IntEREst, blue = KMA). Centroids and whiskers indicate the measure medians and interquartile ranges across persistence cutoffs varied from 0.1 to 0.9 at 0.1 intervals (Methods).

**Fig. S12:**
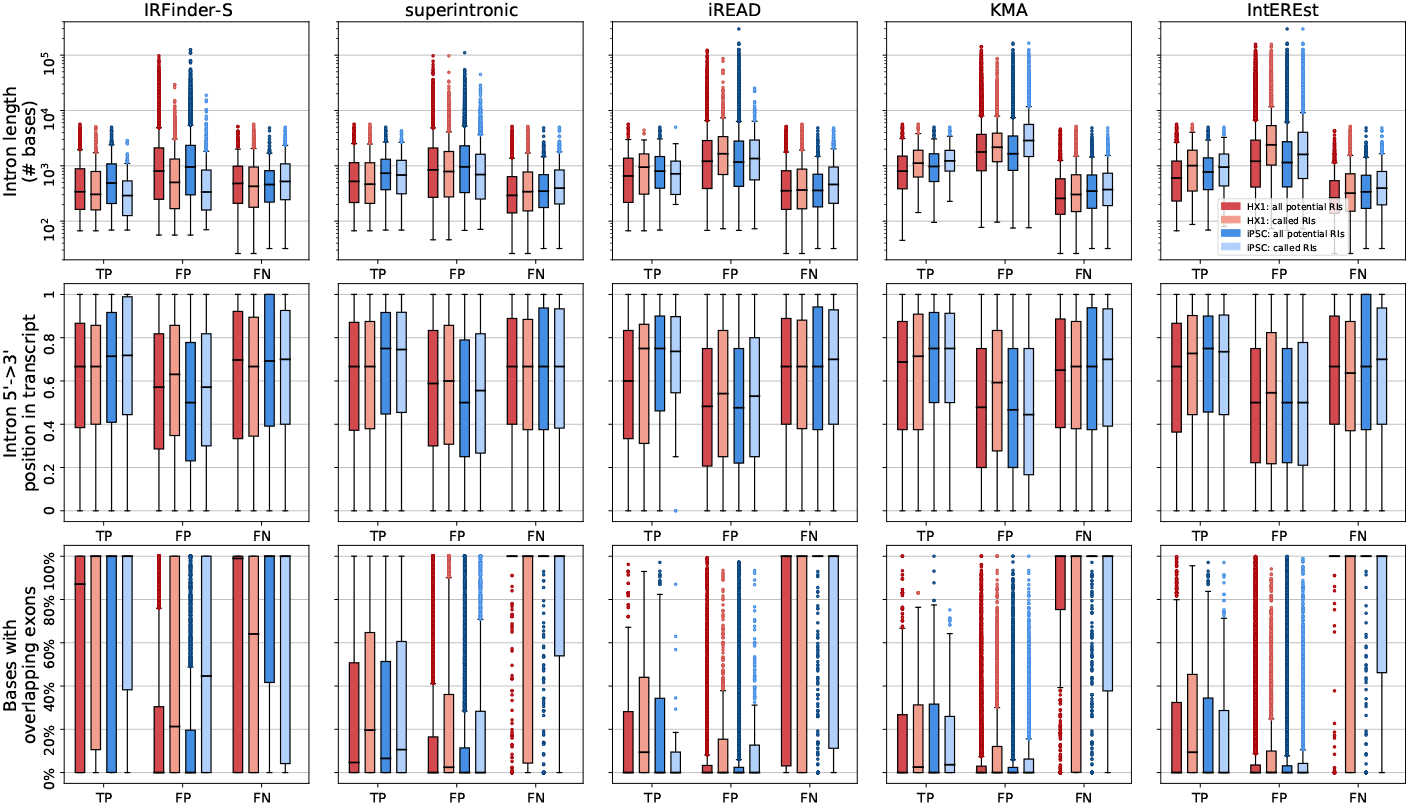
Three target intron properties, length (top), position along the direction of transcription (0 = 5’, 1 = 3’) and % of bases with an overlapping annotated exon vs. TP, FP, and FN calls for HX1 and iPSC via potential RIs (darker) and called RIs (lighter), at long read persistence of 0.1 for 5 short read tools.

**Fig. S13:**
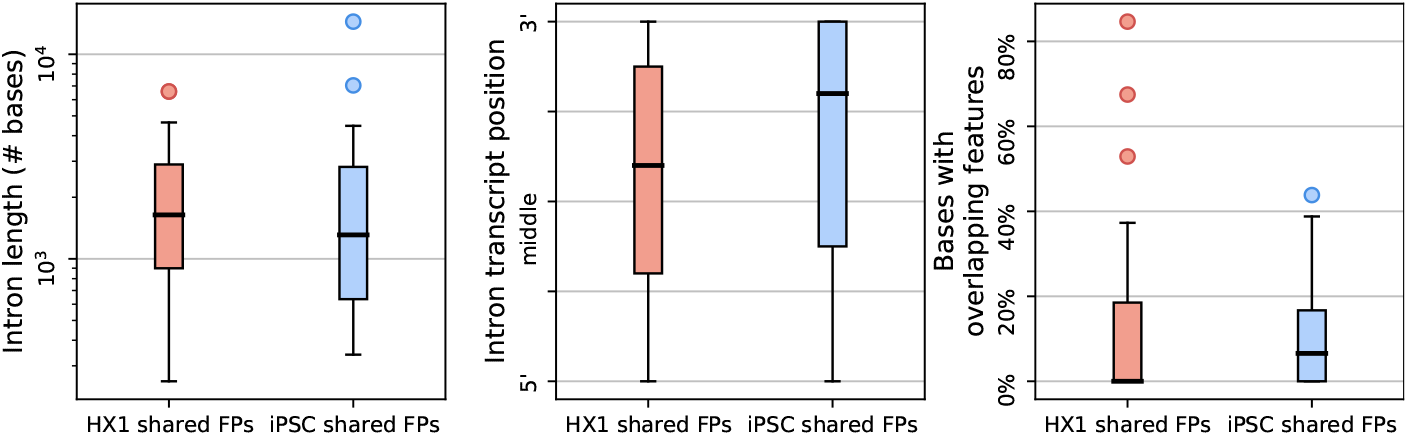
Distribution of intron properties for shared false positive introns (FPs) called across all five short-read detection tools. Properties are, left to right, intron length in # of bases (log scale), transcript position, and % of bases with overlapping exons, for, in each panel, HX1 (left, red) and iPSC (right, blue).

**Fig. S14:**
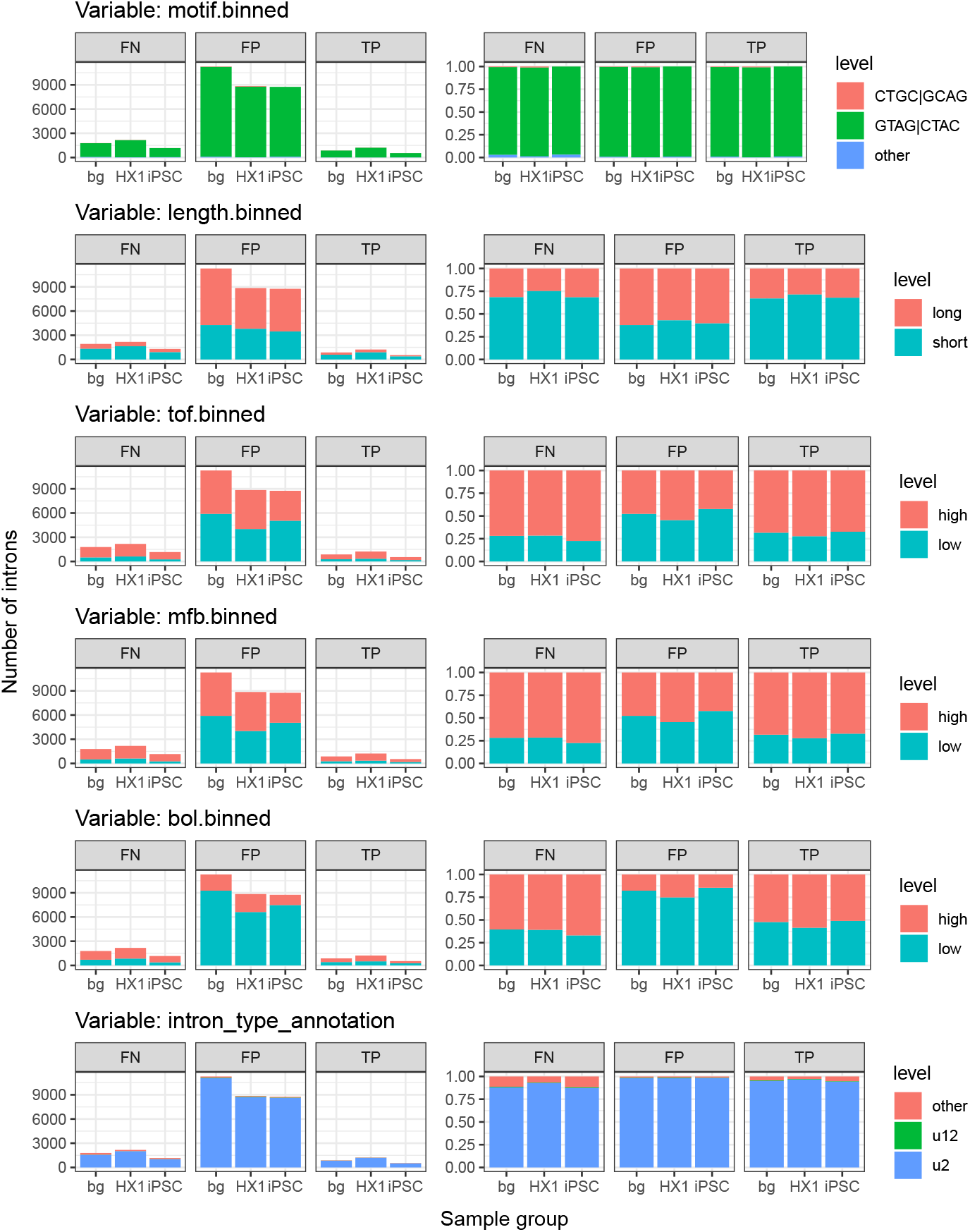
Distributions of binned intron properties. Barplots of intron counts (left column) and percentages (right column) across unique levels (fill colors indicated in legends) for binned intron properties (plot titles). Results were binned by sample group types (columns, either HX1, iPSC, or the background of all unique introns) and intron 4+ truth metric categories TP, FP, and FN (ribbon labels, e.g. intron was TP in at least 4 tools for iPSC, etc.). Qualitative properties were binned by the top three most frequent levels (e.g. “intron type annotation” and “motif binned”), and quantitative properties were binned using the 50th quantile cutoff (e.g. “length”, total overlapping features or “tof,” max features per base or “mfb,” and bases overlapped or “bol”).

**Fig. S15:**
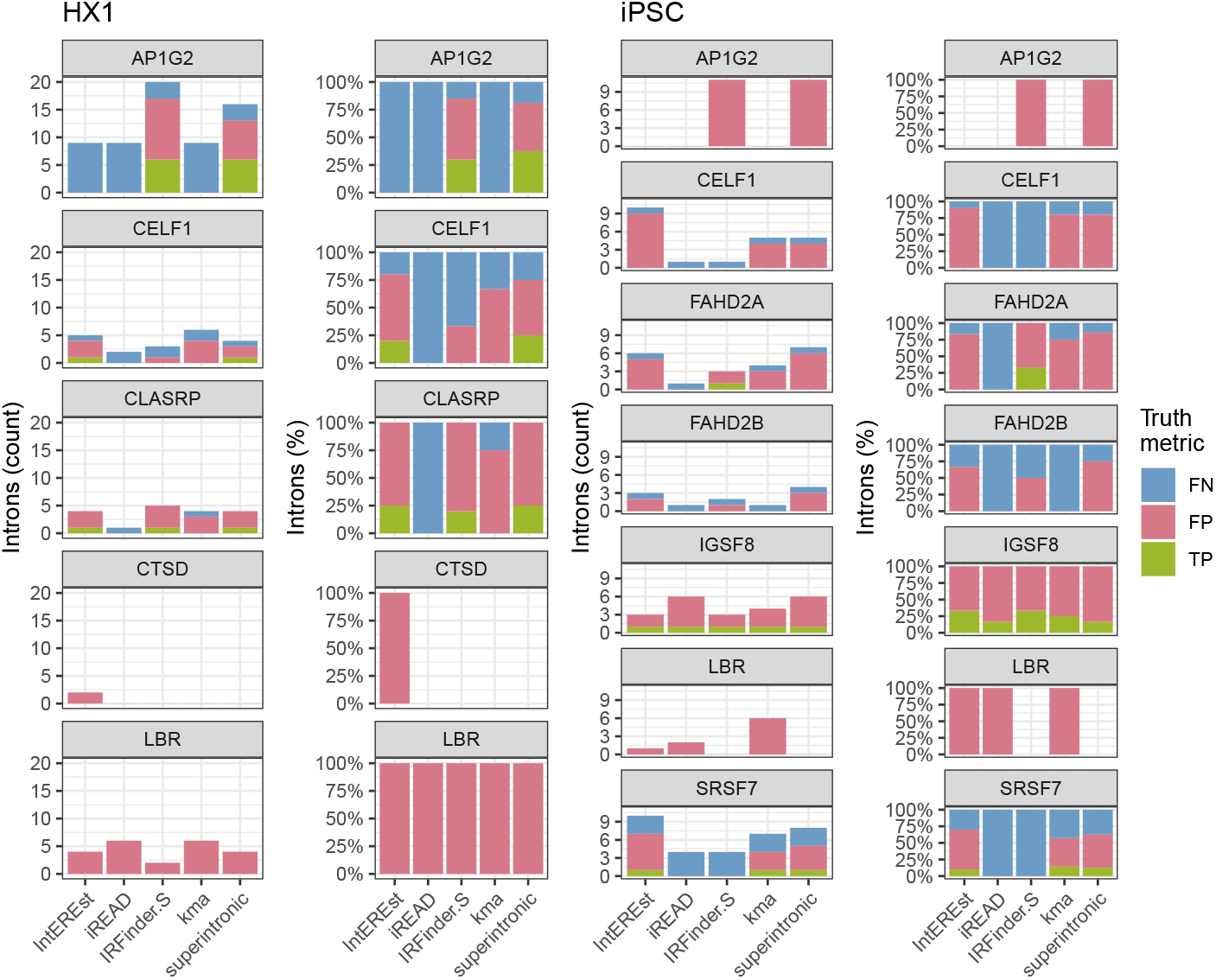
Intron abundance by truth category across genes with validated RIs. Barplots show intron counts and percentages (y-axes) grouped by short-read tool (x-axes), gene (titles), for samples HX1 (left two plot columns) and iPSC (right two plot columns). Bar color fills indicate the short-read tool-specific truth category (green = TP, pink = FP, blue = FN).

**Fig. S16:**
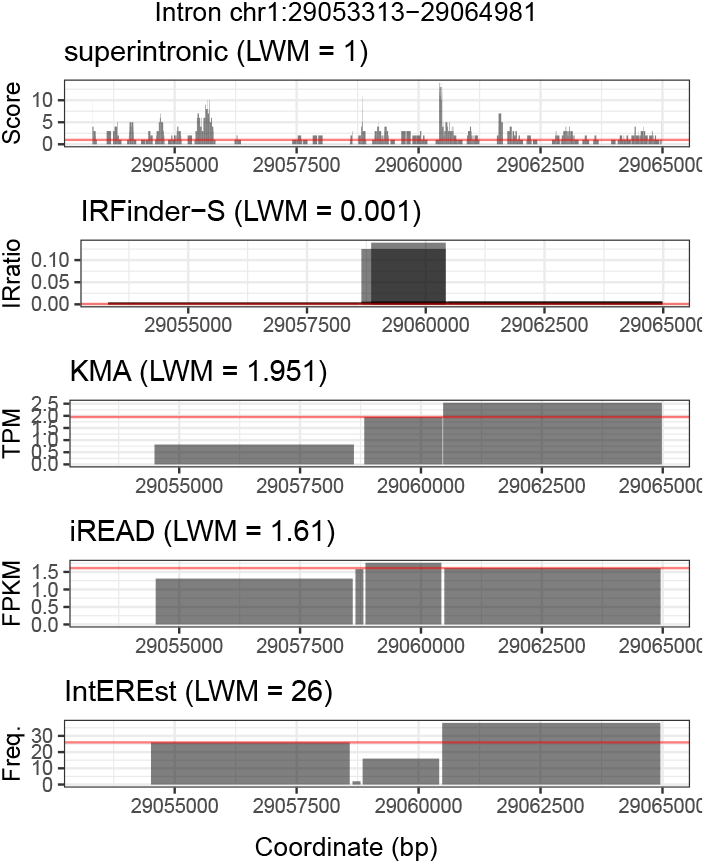
Example length-weighted median expression (LWM) at intron *chr1* :29053313-29064981 (Methods). Intron expression (y-axes) is shown for genomic coordinates (x-axes), where expressed regions are represented by semi-transparent black rectangles which overlap the target intron. Results are grouped by each of the five short-read tools studied, and the LWM value calculated for each tool is shown in the plot titles and horizontal red lines.

**Fig. S17:**
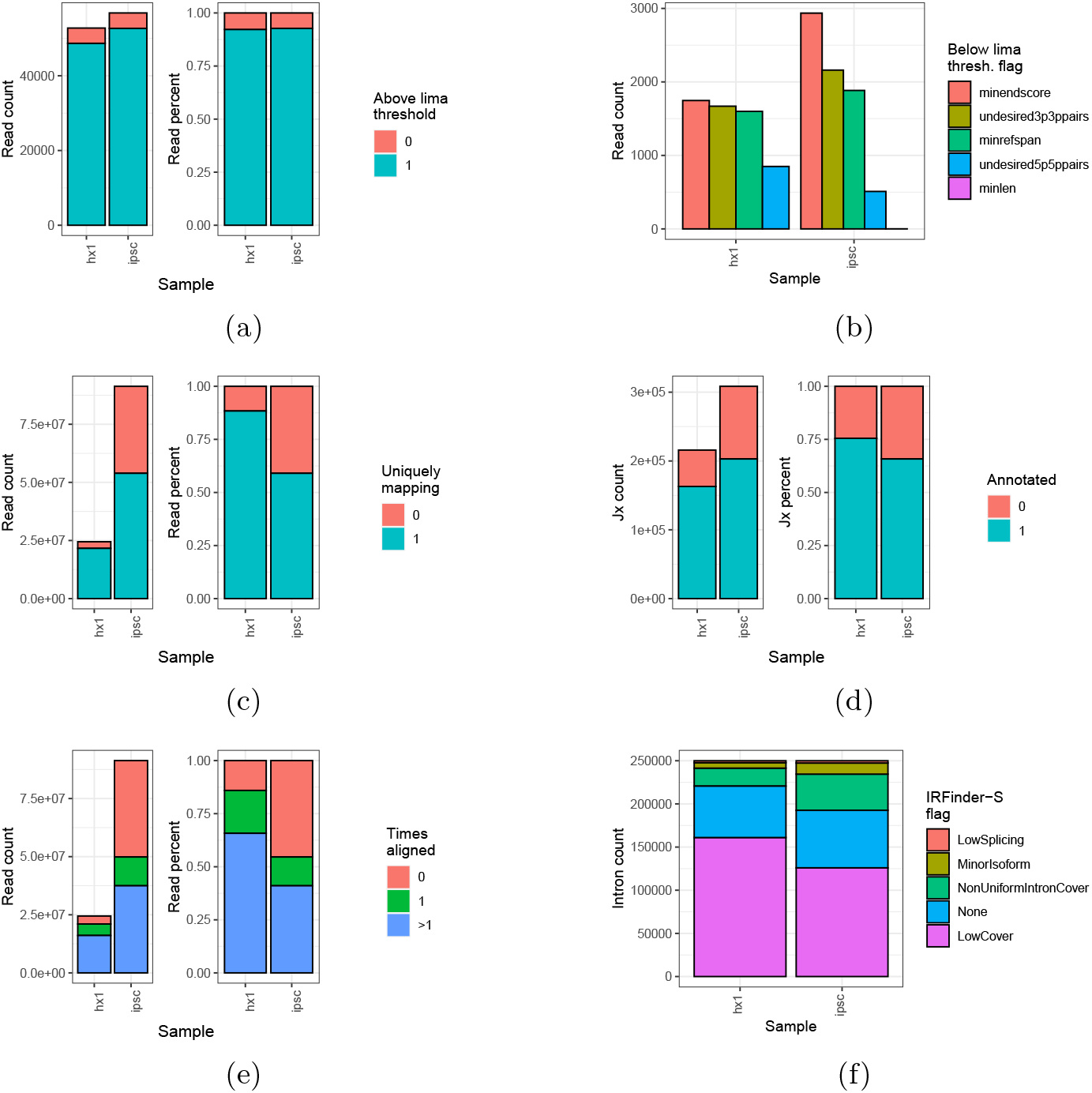
Processing and alignment quality control. Results (y-axes, fill colors) across long-read (A-B) and short-read (C-F) data runs for the samples HX1 and iPSC (x-axes) as follows: (a) LIMA quality among long-reads. Barplot y-axes quantify long-read counts (left) and percentages (right) relative to the quality threshold (blue = above, pink = below), where medians across all runs are shown for iPSC. (b) LIMA flags among long-reads. Barplot y-axes quantify long-reads, where bar colors and x-axes indicate one of the five quality flags (magenta = below minimum length or “minlen”, blue = undesired 5-prime 5-prime pairs as “undesired5p5ppairs”, green = below reference span as “minrefspan”, yellow = undesired 3-prime 3-prime pairs as “undesired3p3ppairs”, and pink = below minimum end score as “minendscore”). (c) Unique mapping among STAR-aligned short-reads. Barplot y-axes quantify short-read counts (left) and percentages (right) by mappability (blue/1 = uniquely mapping, pink/0 = not uniquely mapping). (d) Annotation among STAR-aligned short-reads. Barplot y-axes as in (c) with color indicating annotation (blue/1 = annotated, pink/0 = not annotated). (e) Alignment counts among bowtie2-aligned short-reads. Barplot y-axes as in (c), where bar colors show alignment counts (blue = > 1 times, green = 1 time, pink = 0 times). (f) IRFinder-S flag quantities. Barploy y-axes as in (c), where bar colors show flag (magenta = low coverage as “LowCover”, blue = none, green = non-uniform intron coverage as “NonUniformIntronCover”, yellow = minor isoform presence as “MinorIsoform”, pink = low splicing as “LowSplicing”).

### Supplementary tables

**Table S1:** Description of sequencing data used in this paper.

|  |  |  |
| --- | --- | --- |
| Sample name in paper | iPSC | HX1 |
| Sample type | Induced pluripotent stem cell line | Whole blood (non-cancer) |
| Biosample ID | SAMN07611993 | SAMN04251426 |
| SRA Study ID | SRP098984 | SRP065930 |
| Long read platform | PacBio Iso-Seq RSII | PacBio Iso-Seq RSII |
| Size fractionated | No | Yes |
| Iso-Seq runs | 27 | 46 |
| Aligned long reads | 839,558 | 945,180 |
| Short-read platform | Illumina NextSeq 500 | Illumina HiSeq 2000 |
| Short read runs | 1 | 1 |
| Aligned short reads (% uniquely aligned) | 91,330,785 (59%) | 24,463,210 (88%) |

**Table S2:**
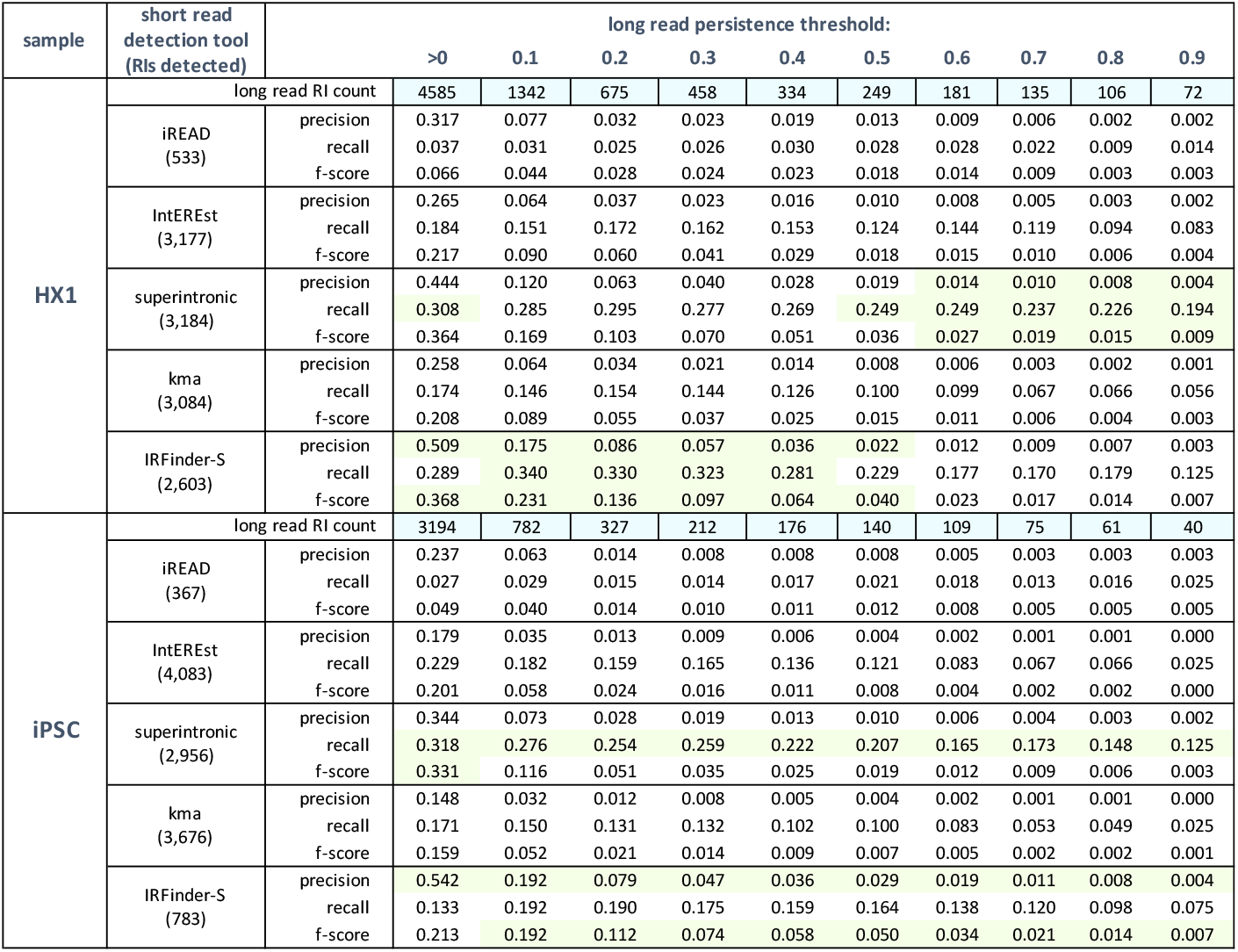
Performance metrics for called RIs across persistence thresholds. Green indicates the highest value per threshold and sample for each metric.

**Table S3:**
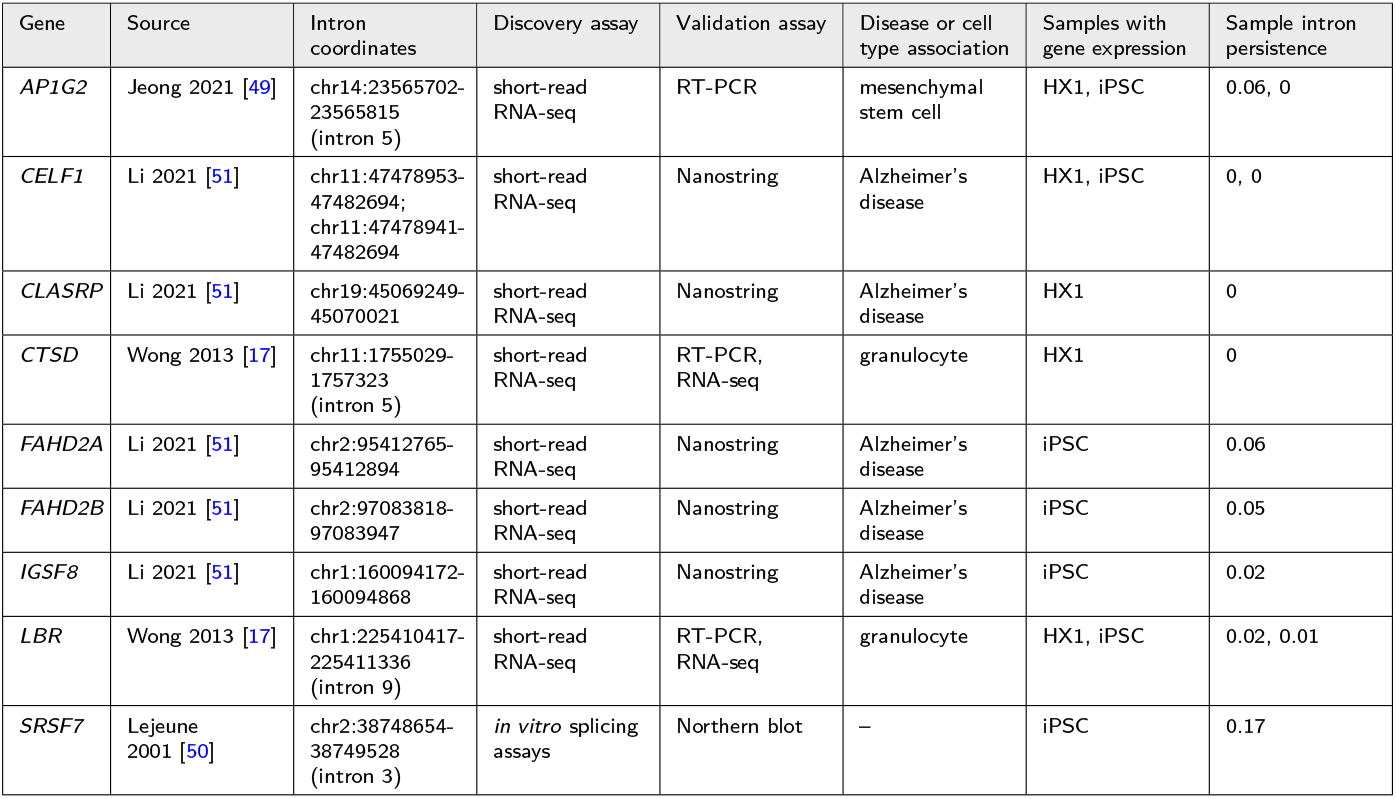
Properties and sources of experimentally validated RIs studied.

| Gene | Source | Intron coordinates | Discovery assay | Validation assay | Disease or cell type association | Samples with gene expression | Sample intron persistence |
| --- | --- | --- | --- | --- | --- | --- | --- |
| *AP1G2* | Jeong 2021 [49] | chr14:23565702-23565815 (intron 5) | short-read RNA-seq | RT-PCR | mesenchymal stem cell | HX1, iPSC | 0.06, 0 |
| *CELF1* | Li 2021 [51] | chr11:47478953-47482694;<br>chr11:47478941-47482694 | short-read RNA-seq | Nanostring | Alzheimer's disease | HX1, iPSC | 0, 0 |
| *CLASRP* | Li 2021 [51] | chr19:45069249-45070021 | short-read RNA-seq | Nanostring | Alzheimer's disease | HX1 | 0 |
| *CTSD* | Wong 2013 [17] | chr11:1755029-1757323 (intron 5) | short-read RNA-seq | RT-PCR, RNA-seq | granulocyte | HX1 | 0 |
| *FAHD2A* | Li 2021 [51] | chr2:95412765-95412894 | short-read RNA-seq | Nanostring | Alzheimer's disease | iPSC | 0.06 |
| *FAHD2B* | Li 2021 [51] | chr2:97083818-97083947 | short-read RNA-seq | Nanostring | Alzheimer's disease | iPSC | 0.05 |
| *IGSF8* | Li 2021 [51] | chr1:160094172-160094868 | short-read RNA-seq | Nanostring | Alzheimer's disease | iPSC | 0.02 |
| *LBR* | Wong 2013 [17] | chr1:225410417-225411336 (intron 9) | short-read RNA-seq | RT-PCR, RNA-seq | granulocyte | HX1, iPSC | 0.02, 0.01 |
| *SRSF7* | Lejeune 2001 [50] | chr2:38748654-38749528 (intron 3) | *in vitro* splicing assays | Northern blot | — | iPSC | 0.17 |

## References

[1] Herzel, Lydia and Ottoz, Diana SM and Alpert, Tara and Neugebauer, Karla M. Splicing and transcription touch base: co-transcriptional spliceosome assembly and function. Nature Reviews Molecular Cell Biology 18 (10), 637–650 (2017) .

[2] Wan, Yihan and Anastasakis, Dimitrios G and Rodriguez, Joseph and Palangat, Murali and Gudla, Prabhakar and Zaki, George and Tandon, Mayank and Pegoraro, Gianluca and Chow, Carson C and Hafner, Markus and others. Dynamic imaging of nascent RNA reveals general principles of transcription dynamics and stochastic splice site selection. Cell 184 (11), 2878–2895 (2021) .

[3] Ameur, Adam and Zaghlool, Ammar and Halvardson, Jonatan and Wetterbom, Anna and Gyllensten, Ulf and Cavelier, Lucia and Feuk, Lars. Total RNA sequencing reveals nascent transcription and widespread co-transcriptional splicing in the human brain. Nature Structural & Molecular Biology 18 (12), 1435–1440 (2011) .

[4] Alpert, Tara and Herzel, Lydia and Neugebauer, Karla M. Perfect timing: Splicing and transcription rates in living cells. Wiley Interdisciplinary Reviews: RNA 8 (2), e1401 (2017) .

[5] Reimer, Kirsten A and Mimoso, Claudia and Adelman, Karen and Neugebauer, Karla M. Rapid and efficient co-transcriptional splicing enhances mammalian gene expression. bioRxiv 2020–02 (2020) .

[6] Girard, Cyrille and Will, Cindy L and Peng, Jianhe and Makarov, Evgeny M and Kastner, Berthold and Lemm, Ira and Urlaub, Henning and Hartmuth, Klaus and Lührmann, Reinhard. Post-transcriptional spliceosomes are retained in nuclear speckles until splicing completion. Nature Communications 3 (1), 1–12 (2012) .

[7] Moyer, Devlin C and Larue, Graham E and Hershberger, Courtney E and Roy, Scott W and Padgett, Richard A. Comprehensive database and evolutionary dynamics of U12-type introns. Nucleic Acids Research 48 (13), 7066–7078 (2020) .

[8] Zhang, Guohong and Taneja, Krishan L and Singer, Robert H and Green, Michael R. Localization of pre-mRNA splicing in mammalian nuclei. Nature 372 (6508), 809–812 (1994) .

[9] Denis, Melvin M and Tolley, Neal D and Bunting, Michaeline and Schwertz, Hansjörg and Jiang, Huimiao and Lindemann, Stephan and Yost, Christian C and Rubner, Frederick J and Albertine, Kurt H and Swoboda, Kathryn J and others. Escaping the nuclear confines: signal-dependent pre-mRNA splicing in anucleate platelets. Cell 122 (3), 379–391 (2005) .

[10] König, Harald and Matter, Nathalie and Bader, Rüdiger and Thiele, Wilko and Müller, Ferenc. Splicing segregation: The minor spliceosome acts outside the nucleus and controls cell proliferation. Cell 131 (4), 718–729 (2007) .

[11] Buckley, Peter T and Khaladkar, Mugdha and Kim, Junhyong and Eberwine, James. Cytoplasmic intron retention, function, splicing, and the sentinel RNA hypothesis. Wiley Interdisciplinary Reviews: RNA 5 (2), 223–230 (2014) .

[12] Uemura, Aya and Oku, Masaya and Mori, Kazutoshi and Yoshida, Hiderou. Unconventional splicing of XBP1 mRNA occurs in the cytoplasm during the mammalian unfolded protein response. Journal of Cell Science 122 (16), 2877–2886 (2009) .

[13] Middleton, Robert and Gao, Dadi and Thomas, Aubin and Singh, Babita and Au, Amy and Wong, Justin JL and Bomane, Alexandra and Cosson, Bertrand and Eyras, Eduardo and Rasko, John EJ and others. IRFinder: assessing the impact of intron retention on mammalian gene expression. Genome Biology 18 (1), 1–11 (2017) .

[14] Yap, Karen and Lim, Zhao Qin and Khandelia, Piyush and Friedman, Brad and Makeyev, Eugene V. Coordinated regulation of neuronal mRNA steady-state levels through developmentally controlled intron retention. Genes & Development 26 (11), 1209–1223 (2012) .

[15] Edwards, Christopher R and Ritchie, William and Wong, Justin J-L and Schmitz, Ulf and Middleton, Robert and An, Xiuli and Mohandas, Narla and Rasko, John EJ and Blobel, Gerd A. A dynamic intron retention program in the mammalian megakaryocyte and erythrocyte lineages. Blood, The Journal of the American Society of Hematology 127 (17), e24–e34 (2016) .

[16] Pimentel, Harold and Parra, Marilyn and Gee, Sherry L and Mohandas, Narla and Pachter, Lior and Conboy, John G. A dynamic intron retention program enriched in RNA processing genes regulates gene expression during terminal erythropoiesis. Nucleic Acids Research 44 (2), 838–851 (2016) .

[17] Wong, Justin J-L and Ritchie, William and Ebner, Olivia A and Selbach, Matthias and Wong, Jason WH and Huang, Yizhou and Gao, Dadi and Pinello, Natalia and Gonzalez, Maria and Baidya, Kinsha and others. Orchestrated intron retention regulates normal granulocyte differentiation. Cell 154 (3), 583–595 (2013) .

[18] Lareau, Liana F and Inada, Maki and Green, Richard E and Wengrod, Jordan C and Brenner, Steven E. Unproductive splicing of SR genes associated with highly conserved and ultraconserved DNA elements. Nature 446 (7138), 926–929 (2007) .

[19] Ge, Ying and Porse, Bo T. The functional consequences of intron retention: alternative splicing coupled to NMD as a regulator of gene expression. Bioessays 36 (3), 236–243 (2014) .

[20] Naro, Chiara and Jolly, Ariane and Di Persio, Sara and Bielli, Pamela and Setterblad, Niclas and Alberdi, Antonio J and Vicini, Elena and Geremia, Raffaele and De la Grange, Pierre and Sette, Claudio. An orchestrated intron retention program in meiosis controls timely usage of transcripts during germ cell differentiation. Developmental Cell 41 (1), 82–93 (2017) .

[21] Memon, Danish and Dawson, Keren and Smowton, Christopher SF and Xing, Wei and Dive, Caroline and Miller, Crispin J. Hypoxia-driven splicing into noncoding isoforms regulates the DNA damage response. npj Genomic Medicine 1 (1), 1–7 (2016) .

[22] Mauger, Oriane and Lemoine, Frédéric and Scheiffele, Peter. Targeted intron retention and excision for rapid gene regulation in response to neuronal activity. Neuron 92 (6), 1266–1278 (2016) .

[23] Ni, Ting and Yang, Wenjing and Han, Miao and Zhang, Yubo and Shen, Ting and Nie, Hongbo and Zhou, Zhihui and Dai, Yalei and Yang, Yanqin and Liu, Poching and others. Global intron retention mediated gene regulation during CD4+ T cell activation. Nucleic Acids Research 44 (14), 6817–6829 (2016) .

[24] Boutz, Paul L and Bhutkar, Arjun and Sharp, Phillip A. Detained introns are a novel, widespread class of post-transcriptionally spliced introns. Genes & Development 29 (1), 63–80 (2015) .

[25] Dvinge, Heidi and Bradley, Robert K. Widespread intron retention diversifies most cancer transcriptomes. Genome Medicine 7 (1), 1–13 (2015) .

[26] Zhang, Qu and Li, Hua and Jin, Hong and Tan, Huibiao and Zhang, Jun and Sheng, Sitong. The global landscape of intron retentions in lung adenocarcinoma. BMC Medical Genomics 7 (1), 1–9 (2014) .

[27] Eswaran, Jeyanthy and Horvath, Anelia and Godbole, Sucheta and Reddy, Sirigiri Divijendra and Mudvari, Prakriti and Ohshiro, Kazufumi and Cyanam, Dinesh and Nair, Sujit and Fuqua, Suzanne AW and Polyak, Kornelia and others. RNA sequencing of cancer reveals novel splicing alterations. Scientific Reports 3 (1), 1–12 (2013) .

[28] Monteuuis, Geoffray and Wong, Justin JL and Bailey, Charles G and Schmitz, Ulf and Rasko, John EJ. The changing paradigm of intron retention: Regulation, ramifications and recipes. Nucleic Acids Research 47 (22), 11497–11513 (2019) .

[29] de Lima Morais, David A and Harrison, Paul M. Large-scale evidence for conservation of NMD candidature across mammals. PLoS One 5 (7), e11695 (2010) .

[30] Buckley, Peter T and Lee, Miler T and Sul, Jai-Yoon and Miyashiro, Kevin Y and Bell, Thomas J and Fisher, Stephen A and Kim, Junhyong and Eberwine, James. Cytoplasmic intron sequence-retaining transcripts can be dendritically targeted via ID element retrotransposons. Neuron 69 (5), 877–884 (2011) .

[31] Smart, Alicia C and Margolis, Claire A and Pimentel, Harold and He, Meng Xiao and Miao, Diana and Adeegbe, Dennis and Fugmann, Tim and Wong, Kwok-Kin and Van Allen, Eliezer M. Intron retention is a source of neoepitopes in cancer. Nature Biotechnology 36 (11), 1056–1058 (2018) .

[32] Kahles, André and Lehmann, Kjong-Van and Toussaint, Nora C and Hüser, Matthias and Stark, Stefan G and Sachsenberg, Timo and Stegle, Oliver and Kohlbacher, Oliver and Sander, Chris and Caesar-Johnson, Samantha J and others. Comprehensive analysis of alternative splicing across tumors from 8,705 patients. Cancer Cell 34 (2), 211–224 (2018) .

[33] Trincado, Juan L and Reixachs-Sole, Marina and Pérez-Granado, Judith and Fugmann, Tim and Sanz, Ferran and Yokota, Jun and Eyras, Eduardo. ISOTOPE: ISOform-guided prediction of epiTOPEs in cancer. PLoS Computational Biology 17 (9), e1009411 (2021) .

[34] Dong, Chuanpeng and Cesarano, Annamaria and Bombaci, Giuseppe and Reiter, Jill L and Yu, Christina Y and Wang, Yue and Jiang, Zhaoyang and Zaid, Mohammad Abu and Huang, Kun and Lu, Xiongbin and others. Intron retention-induced neoantigen load correlates with unfavorable prognosis in multiple myeloma. Oncogene 40 (42), 6130–6138 (2021) .

[35] Dong, Chuanpeng and Reiter, Jill L and Dong, Edward and Wang, Yue and Lee, Kelvin P and Lu, Xiongbin and Liu, Yunlong. Intron-Retention neoantigen load predicts favorable prognosis in pancreatic cancer. JCO Clinical Cancer Informatics 6, e2100124 (2022) .

[36] Szabo, Linda and Salzman, Julia. Detecting circular RNAs: Bioinformatic and experimental challenges. Nature Reviews Genetics 17 (11), 679–692 (2016) .

[37] Pimentel, Harold and Conboy, John G and Pachter, Lior. Keep me around: Intron retention detection and analysis. arXiv preprint arXiv:1510.00696 (2015) .

[38] Oghabian, Ali and Greco, Dario and Frilander, Mikko J. IntEREst: Intron-exon retention estimator. BMC Bioinformatics 19 (1), 1–10 (2018) .

[39] Li, Hong-Dong and Funk, Cory C and Price, Nathan D. iREAD: a tool for intron retention detection from RNA-seq data. BMC Genomics 21 (1), 1–11 (2020) .

[40] Lee, Stuart and Zhang, Albert Y and Su, Shian and Ng, Ashley P and Holik, Aliaksei Z and Asselin-Labat, Marie-Liesse and Ritchie, Matthew E and Law, Charity W. Covering all your bases: Incorporating intron signal from RNA-seq data. NAR Genomics and Bioinformatics 2 (3), lqaa073 (2020) .

[41] Lorenzi, Claudio and Barriere, Sylvain and Arnold, Katharina and Luco, Reini F and Oldfield, Andrew J and Ritchie, William. IRFinder-S: A comprehensive suite to discover and explore intron retention. Genome Biology 22 (1), 1–13 (2021) .

[42] Tilgner, Hagen and Knowles, David G and Johnson, Rory and Davis, Carrie A and Chakrabortty, Sudipto and Djebali, Sarah and Curado, João and Snyder, Michael and Gingeras, Thomas R and Guigó, Roderic. Deep sequencing of subcellular RNA fractions shows splicing to be predominantly co-transcriptional in the human genome but inefficient for lncRNAs. Genome Research 22 (9), 1616–1625 (2012) .

[43] Braunschweig, Ulrich and Barbosa-Morais, Nuno L and Pan, Qun and Nachman, Emil N and Alipanahi, Babak and Gonatopoulos-Pournatzis, Thomas and Frey, Brendan and Irimia, Manuel and Blencowe, Benjamin J. Widespread intron retention in mammals functionally tunes transcriptomes. Genome Research 24 (11), 1774–1786 (2014) .

[44] Shi, Lingling and Guo, Yunfei and Dong, Chengliang and Huddleston, John and Yang, Hui and Han, Xiaolu and Fu, Aisi and Li, Quan and Li, Na and Gong, Siyi and others. Long-read sequencing and de novo assembly of a Chinese genome. Nature Communications 7 (1), 1–10 (2016) .

[45] Kronenberg, Zev N and Fiddes, Ian T and Gordon, David and Murali, Shwetha and Cantsilieris, Stuart and Meyerson, Olivia S and Underwood, Jason G and Nelson, Bradley J and Chaisson, Mark JP and Dougherty, Max L and others. High-resolution comparative analysis of great ape genomes. Science 360 (6393), eaar6343 (2018) .

[46] Glažar, Petar and Papavasileiou, Panagiotis and Rajewsky, Nikolaus. circBase: A database for circular RNAs. RNA 20 (11), 1666–1670 (2014) .

[47] User bulletin: Guidelines for preparing cDNA libraries for isoform sequencing (Iso-Seq (TM)). https://www.pacb.com/wp-content/uploads/2015/09/User-Bulletin-Guidelines-for-Preparing-cDNA-Libraries-for-Isoform-Sequencing-Iso-Seq.pdf. Accessed: 2022-03-02.

[48] Khrameeva, Ekaterina E and Gelfand, Mikhail S. Biases in read coverage demonstrated by interlaboratory and interplatform comparison of 117 mRNA and genome sequencing experiments. BMC Bioinformatics 13 (6), 1–7 (2012) .

[49] Jeong, Ji-Eun and Seol, Binna and Kim, Han-Seop and Kim, Jae-Yun and Cho, Yee-Sook. Exploration of alternative splicing events in mesenchymal stem cells from human induced pluripotent stem cells. Genes 12 (5), 737xs (2021) .

[50] Lejeune, Fabrice and Cavaloc, Yvon and Stevenin, James. Alternative splicing of intron 3 of the serine/arginine-rich protein 9G8 gene: Identification of flanking exonic splicing enhancers and involvement of 9G8 as a trans-acting factor. Journal of Biological Chemistry 276 (11), 7850–7858 (2001) .

[51] Li, Hong-Dong and Funk, Cory C and McFarland, Karen and Dammer, Eric B and Allen, Mariet and Carrasquillo, Minerva M and Levites, Yona and Chakrabarty, Paramita and Burgess, Jeremy D and Wang, Xue and others. Integrative functional genomic analysis of intron retention in human and mouse brain with Alzheimer’s disease. Alzheimer’s & Dementia 17 (6), 984–1004 (2021) .

[52] Königs, Vanessa and de Oliveira Freitas Machado, Camila and Arnold, Benjamin and Blümel, Nicole and Solovyeva, Anfisa and Löbbert, Sinah and Schafranek, Michal and Ruiz De Los Mozos, Igor and Wittig, Ilka and McNicoll, Francois and others. SRSF7 maintains its homeostasis through the expression of split-ORFs and nuclear body assembly. Nature Structural & Molecular Biology 27 (3), 260–273 (2020) .

[53] Heinicke, Laurie A and Nabet, Behnam and Shen, Shihao and Jiang, Peng and van Zalen, Sebastiaan and Cieply, Benjamin and Russell, J Eric and Xing, Yi and Carstens, Russ P. The RNA binding protein RBM38 (RNPC1) regulates splicing during late erythroid differentiation. PloS ONE 8 (10), e78031 (2013) .

[54] Bhatt, Dev M and Pandya-Jones, Amy and Tong, Ann-Jay and Barozzi, Iros and Lissner, Michelle M and Natoli, Gioacchino and Black, Douglas L and Smale, Stephen T. Transcript dynamics of proinflammatory genes revealed by sequence analysis of subcellular RNA fractions. Cell 150 (2), 279–290 (2012) .

[55] Schmitz, Ulf and Pinello, Natalia and Jia, Fangzhi and Alasmari, Sultan and Ritchie, William and Keightley, Maria-Cristina and Shini, Shaniko and Lieschke, Graham J and Wong, Justin JL and Rasko, John EJ. Intron retention enhances gene regulatory complexity in vertebrates. Genome Biology 18 (1), 1–15 (2017) .

[56] Sakabe, Noboru Jo and De Souza, Sandro José. Sequence features responsible for intron retention in human. BMC Genomics 8 (1), 1–14 (2007) .

[57] Li, Heng and Handsaker, Bob and Wysoker, Alec and Fennell, Tim and Ruan, Jue and Homer, Nils and Marth, Gabor and Abecasis, Goncalo and Durbin, Richard. The sequence alignment/map format and SAMtools. Bioinformatics 25 (16), 2078–2079 (2009) .

[58] Bonfield, James K and Marshall, John and Danecek, Petr and Li, Heng and Ohan, Valeriu and Whitwham, Andrew and Keane, Thomas and Davies, Robert M. HTSlib: C library for reading/writing high-throughput sequencing data. Gigascience 10 (2), giab007 (2021) .

[59] Dobin, Alexander and Davis, Carrie A and Schlesinger, Felix and Drenkow, Jorg and Zaleski, Chris and Jha, Sonali and Batut, Philippe and Chaisson, Mark and Gingeras, Thomas R. STAR: Ultrafast universal RNA-seq aligner. Bioinformatics 29 (1), 15–21 (2013) .

[60] Frankish, Adam and Diekhans, Mark and Ferreira, Anne-Maud and Johnson, Rory and Jungreis, Irwin and Loveland, Jane and Mudge, Jonathan M and Sisu, Cristina and Wright, James and Armstrong, Joel and others. GENCODE reference annotation for the human and mouse genomes. Nucleic Acids Research 47 (D1), D766–D773 (2019) .

[61] Langmead, Ben and Salzberg, Steven L. Fast gapped-read alignment with Bowtie 2. Nature Methods 9 (4), 357–359 (2012) .

[62] pachterlab/kma: Keep Me Around: Intron retention detection. https://github.com/pachterlab/kma. Accessed: 2022-03-02.

[63] Roberts, Adam and Pachter, Lior. Streaming fragment assignment for real-time analysis of sequencing experiments. Nature Methods 10 (1), 71–73 (2013) .

[64] Roberts, Adam and Trapnell, Cole and Donaghey, Julie and Rinn, John L and Pachter, Lior. Improving RNA-Seq expression estimates by correcting for fragment bias. Genome Biology 12 (3), 1–14 (2011) .

[65] Li, Hong-Dong. GTFtools: A Python package for analyzing various modes of gene models. bioRxiv 263517 (2018) .

[66] Fleiss, Joseph L. Measuring nominal scale agreement among many raters. Psychological Bulletin 76 (5), 378 (1971) .

[67] irr: Various Coefficients of Interrater Reliability and Agreement. https://CRAN.R-project.org/package=irr. Accessed: 2022-03-08.

[68] Dumbović, Gabrijela and Braunschweig, Ulrich and Langner, Heera K and Smallegan, Michael and Biayna, Josep and Hass, Evan P and Jastrzebska, Katarzyna and Blencowe, Benjamin and Cech, Thomas R and Caruthers, Marvin H and others. Nuclear compartmentalization of TERT mRNA and TUG1 lncRNA is driven by intron retention. Nature Communications 12 (1), 1–19 (2021) .

[69] Inoue, Daichi and Polaski, Jacob T and Taylor, Justin and Castel, Pau and Chen, Sisi and Kobayashi, Susumu and Hogg, Simon J and Hayashi, Yasutaka and Pineda, Jose Mario Bello and El Marabti, Ettaib and others. Minor intron retention drives clonal hematopoietic disorders and diverse cancer predisposition. Nature Genetics 53 (5), 707–718 (2021) .

[70] Bell, Thomas J and Miyashiro, Kevin Y and Sul, Jai-Yoon and McCullough, Ronald and Buckley, Peter T and Jochems, Jeanine and Meaney, David F and Haydon, Phil and Cantor, Charles and Parsons, Thomas D and others. Cytoplasmic BKCa channel intron-containing mRNAs contribute to the intrinsic excitability of hippocampal neurons. Proceedings of the National Academy of Sciences 105 (6), 1901–1906 (2008) .

[71] Bell, Thomas J and Miyashiro, Kevin Y and Sul, Jai-Yoon and Buckley, Peter T and Lee, Miler T and McCullough, Ron and Jochems, Jeanine and Kim, Junhyong and Cantor, Charles R and Parsons, Thomas D and others. Intron retention facilitates splice variant diversity in calcium-activated big potassium channel populations. Proceedings of the National Academy of Sciences 107 (49), 21152–21157 (2010) .

[72] Chen, Zhe and Gore, Bryan B and Long, Hua and Ma, Le and Tessier-Lavigne, Marc. Alternative splicing of the Robo3 axon guidance receptor governs the midline switch from attraction to repulsion. Neuron 58 (3), 325–332 (2008) .

[73] Zhong, Xiaoli and Liu, Jinrong R and Kyle, John W and Hanck, Dorothy A and Agnew, William S. A profile of alternative RNA splicing and transcript variation of CACNA1H, a human T-channel gene candidate for idiopathic generalized epilepsies. Human Molecular Genetics 15 (9), 1497–1512 (2006) .

[74] Mansilla, Alicia and López-Sánchez, Carmen and De la Rosa, Enrique J and García-Martínez, Virginio and Martínez-Salas, Encarna and de Pablo, Flora and Hernández-Sánchez, Catalina. Developmental regulation of a proinsulin messenger RNA generated by intron retention. EMBO Reports 6 (12), 1182–1187 (2005) .

[75] Forrest, Scott T and Barringhaus, Kurt G and Perlegas, Demetra and Hammarskjold, Marie-Louise and McNamara, Coleen A. Intron retention generates a novel Id3 isoform that inhibits vascular lesion formation. Journal of Biological Chemistry 279 (31), 32897–32903 (2004) .

